# Persistent light-induced reduction of neuronal excitability in cortical neurons

**DOI:** 10.1101/2025.04.25.650615

**Authors:** Anistasha Lightning, Jean-Francois Ghersi-Egea, Federico Di Rocco, Marc Guenot, Nicola Kuczewski

## Abstract

Visible light is widely used in neuroscience, yet its direct effects on neuronal activity in the absence of optogenetic manipulation remain incompletely understood. Here, we investigated whether light stimulation can induce sustained changes in neuronal excitability. Using electrophysiological recordings in brain slices, we show that repeated pulses of blue light (5 s, 430–495 nm, 19 mW) produce a robust and persistent reduction in evoked firing activity in cortical neurons from both male and female mice, with an average decrease of ∼60% relative to baseline. This inhibitory effect persisted for more than 20 minutes following stimulation and was associated with changes in both passive membrane properties and active ion channel conductance. To assess the potential translational relevance of visible-light stimulation as a future therapeutic approach for brain pathologies, we examined its effects in human cortical neurons where the, responses were more heterogeneous. While a subset of neurons exhibited similar inhibitory effects, others showed increased excitability, with this response occurring more frequently in neurons from female patients, suggesting a potential sex-dependent effect, which ultimately remains exploratory in the present work. In addition, a transient depolarizing response to light was observed in a minority of human neurons but not in mice. Because the contribution of light-induced heating was not experimentally isolated, the relative roles of photochemical and thermal mechanisms in these effects remain unresolved. These findings indicate that visible light exposure, independently of exogenous opsins, can induce long-lasting modulation of neuronal activity without evidence of acute cytotoxicity under the conditions tested. This raises the possibility that visible light may provide a previously underappreciated, opsin-independent mechanism for modulating neuronal activity, with potential relevance for disorders characterized by neuronal hyperexcitability. We outline a framework for future investigations, including validation in human systems, in vivo studies, optimization of stimulation parameters, and assessment of therapeutic potential in pathological models.

## Introduction

Although pharmacological treatments are the main approach to neuronal hyperexcitability, they have several limitations, including poor regional specificity, difficulty crossing the blood–brain barrier, peripheral side effects, and the risk of tolerance and dependence with prolonged use ^1,2^. These limitations have prompted the development of targeted, non-pharmacological approaches, such as Deep Brain Stimulation (DBS)^3^, Transcranial Magnetic Stimulation^4^, and ultrasound neuromodulation^5^. These techniques are widely used in research, and DBS is an established treatment for selected patients with advanced Parkinson’s disease^6^. Another approach is light-based neuromodulation. Near-infrared photobiomodulation may influence neuronal metabolism by reducing oxidative stress, modulating inflammation, and promoting neurogenesis^7,8^. More recently, studies have shown that visible light (yellow-to-blue wavelengths) can also reduce neuronal firing across various central nervous system cell types^9–11^.However, the light effects are predominantly transient, with neuronal activity recovering shortly after illumination ceases. Short lasting inhibition appears to involve a light-induced hyperpolarizing current and changes in action potential (AP) amplitude and latency^9,12^ Although the underlying mechanisms remain unclear, evidence suggests that light-induced heating alters membrane channel gating and/or permeability, affecting neuronal activity ^9,10,12,13^. Inhibition increases with light power and stimulation duration^9,13,14^. Whether visible light can induce persistent changes in neuronal excitability that outlast the stimulation period remains largely unexplored.Preliminary findings further suggest that repeated 5–10 s blue light pulses induce sustained firing inhibition in olfactory bulb mitral cells, persisting for several minutes after stimulation ^9^. These observations raised the possibility that appropriately designed visible-light stimulation could produce persistent, rather than solely acute, modulation of neuronal excitability, suggesting that blue light stimulation could be used as a therapeutical tool for treating the neuronal hyperexcitability associated with several neurological disorders in humans in the absence of the expression of exogenous opsin in the brain. The aim of the present study was therefore to determine whether repeated blue-light stimulation can induce persistent changes in neuronal excitability that outlast the illumination period, and to characterize these effects in both mouse and human cortical neurons.

## Material and methods

### Animals

Male and female C57BL/6J mice (60–90 days old; Janvier Laboratories, France) were used. Animals were euthanized solely for tissue collection, with no experimental procedures performed on live animals; therefore, no specific animal experiment licence or protocol number applied. All procedures complied with EU Directive 2010/63/EU and institutional animal welfare requirements. Mice were housed in groups of up to six under a 12-h light/dark cycle at 22°C, with food and water ad libitum.

### Human Tissue

To investigate cross-species similarities and potential clinical relevance, human cortical tissue was obtained from residual surgical specimens collected during procedures at two hospitals in Bron, France. Samples from patients of both sexes (aged 3–54 years) undergoing surgery for drug-resistant epilepsy or brain tumors were used in accordance with French regulations for human biological samples (CODECOH declaration No. DC 2019-3668). Informed consent was obtained from all patients or their legal guardians, as applicable.

### Optical stimulation

Single-photon blue light stimulation (peak wavelength: 470 nm; emission spectrum: 430–495 nm) was delivered using a Dual Port OptoLED system (CAIRN, UK) at near-maximum output power (19 mW), measured through a 40× objective. Power loss due to ACSF at the working distance (2.5 mm) was estimated at 13%, based on measurements with and without ACSF. The resulting average power density was estimated at ∼2.4 mW/mm² over an illumination area of ∼7 mm². As each measurement was performed once, no variability estimate was available. Stimulation duration was set to 5 s to reproduce the sustained firing inhibition previously reported in olfactory bulb mitral cells^9^.

### Slice preparation of mouse cortical neurons

Mice were anesthetized with intraperitoneal ketamine (50 mg/ml) and decapitated. The head was rapidly immersed in ice-cold (2–4°C) cutting artificial cerebrospinal fluid (ACSF; 125 mM NaCl, 4 mM KCl, 25 mM NaHCO₃, 0.5 mM CaCl₂, 1.25 mM NaH₂PO₄, 7 mM MgCl₂, and 5.5 mM glucose; pH 7.4), oxygenated with 95% O₂/5% CO₂. Osmolarity was adjusted to 300–320 mOsm with sucrose. After brain extraction, the cerebellum, midbrain, olfactory bulb, and approximately 1–2 mm of anterior cortex were removed with a razor blade to obtain a flat surface for vibratome mounting. Coronal cortical slices (400 µm) were prepared using a Leica vibratome and bisected along the corpus callosum. Slices were then incubated in a recovery chamber at 30 ± 1°C in ACSF with the same composition, except for CaCl₂ and MgCl₂ concentrations (1.2 and 0.7 mM, respectively).

### Slice preparation of human neurons

Approximately 0.5–2 cm³ of temporal cortical tissue was collected by the neurosurgeon and immediately transferred to ice-cold (2–4°C), oxygenated cutting ACSF, as described for mouse tissue. During transport, the solution was continuously oxygenated with 95% O₂/5% CO₂ using a portable oxygen tank. Upon arrival, the tissue was trimmed to fit the vibratome chamber, with slicing performed 10–20 min after resection. Coronal slices (400 µm) were prepared, retaining both gray and white matter when possible to facilitate orientation under the microscope. Visual inspection revealed no obvious tissue degradation following transport.

### Determination of light-induced tissue temperature change

LED-induced temperature changes were measured using a ThermoClamp-1 device (Atomate Scientific; accuracy <1 °C, resolution 0.1 °C). The probe was placed on cortical tissue beneath the microscope objective at the same distance as in electrophysiological experiments. Temperature was recorded during 5-s stimulation with 19 mW continuous blue light.

### Electrophysiological recordings

Slices were perfused with oxygenated ACSF (2–4 ml/min) at 36 ± 1°C in a recording chamber mounted on an upright microscope (Zeiss Axioscope, ×40 Plan-Apochromat objective). Data were acquired at 25 kHz using an RK 400 amplifier (BioLogic), a 12-bit Digidata 1440A converter, and PClamp 10 software (Axon Instruments). Whole-cell recordings were obtained from cortical neurons across different regions and layers using borosilicate pipettes (5–9 MΩ) filled with (in mM): 126 K-gluconate, 5 KCl, 10 HEPES, 1 EGTA, 1 MgCl₂, 2 ATP-Na₂, 0.3 GTP-Na₃, and 10 phosphocreatine (pH 7.3, 290 mOsm). Excitatory glutamatergic neurons were identified by their pyramidal morphology, ISI coefficient of variation >0.3, firing adaptation ratio (last/first ISI) >1.5, and AP duration (FWHM) >0.5 ms. Neurons were sampled across different cortical layers to accommodate variability in the cortical areas available from human tissue and to increase the likelihood of obtaining suitable recordings despite variations in slice quality. This approach allowed cross-species comparisons while reducing the number of animals required.

### Determination of the light effect on the evoked firing rate of cortical neurons

This protocol was applied to mouse and human neurons in current-clamp mode at resting membrane potential, except for one neuron in which a small hyperpolarizing current suppressed spontaneous activity. Recordings consisted of 38-s sweeps (Supplementary Fig. 1), each including a 1-s, −0.1 nA hyperpolarizing step to assess membrane resistance (Rm), followed by two 5-s depolarizing steps separated by 15 s. After nine control sweeps (6 min), six or ten sweeps were performed with 5-s blue-light stimulation (19 mW) during the first depolarizing step (StimLED), but not the second (StimCtr). An additional 1-s light pulse was delivered 4 s after the second step to assess resting membrane potential changes. Post-light recordings continued for 20–42 min. A custom Python script (https://osf.io/unj9k/files/whs5m/) quantified firing-rate changes relative to the median control rate. Light-induced Vm changes were calculated as the difference between mean Vm during the light pulse and the preceding 1 s. Rm was calculated using Ohm’s law (Rm = ΔV/ΔI), with ΔV defined as the difference between Vm during the final 100 ms of the hyperpolarizing step and the preceding 200 ms.

### Determination of light effect on the currents participating in the action potential generation

Mouse cortical neurons were recorded in voltage-clamp mode at −70 mV (Supplementary Fig. 2). Dynamic AP clamp used a voltage command reproducing a 78-mV action potential recorded in current-clamp mode^12^. Ten preceding waveforms scaled by −0.1 (−7.8 to +0.4 mV) elicited passive currents, which were subtracted from total currents to isolate active inward Na⁺ and outward K⁺ currents. Recordings comprised 13 control sweeps, six LED-stimulation sweeps, and 25 min of post-stimulation monitoring. To correct for access resistance (Rₐ) variations, each current trace was multiplied by the ratio of Rₐ in the first sweep to that in the corresponding sweep. The effectiveness of this correction in compensating for changes in current amplitude and kinetics illustrated in Supplementary Fig. 2D**, a**lthough **it** may vary across recordings. A 1-s, −5-mV hyperpolarizing step preceding each waveform assessed membrane resistance (Rₘ = ΔV/ΔI), with ΔI calculated from median currents during the final 100 ms of the step and the preceding 200 ms. Rₐ was calculated as ΔV/ΔIₐ, where ΔIₐ represents the difference between peak transient and median baseline currents. Analyses used custom Python scripts (https://osf.io/unj9k/files/j7u4k).

### Superoxide quantification

Superoxide anion produced by brain slices and released in the medium was quantified using acetylated cytochrome c as a trapping agent, as previously described in detail^15^. Acetylated cytochrome c is specifically and stoichiometrically reduced by superoxide anions. Briefly, Acetylated cytochrome c (Sigma C-4186, 99% oxidized form) was added in the medium at a final concentration of 60 µM. At different time interval the medium was sampled and its absorbance spectrum recorded in a Cary 100 Spectrophotometer (Varian). The % of reduced acetylated cytochrome c was calculated from the corresponding second derivative spectrum as detailed in supplementary figure 3.

### Statistics

Biophysical parameters were compared across pre-light, light, and post-light conditions using repeated-measures ANOVA. Multiple comparisons were adjusted using Holm’s method, either for post hoc pairwise comparisons or for the set of planned contrasts, as appropriate. Pearson’s correlation, 2 × 2 repeated-measures ANOVA (sex × light), and χ² tests were used as appropriate. Significance was set at α = 0.05 evidence for the null hypothesis was assessed using the Bayes factor (BF₀₁). Most analyses were performed at the neuron level. For most of the analysis animal identity was not included as a random effect because the limited number of neurons per animal precluded reliable estimation of between-animal variability. Where appropriate, multilevel models or animal-/subject-level aggregation were used to account for data nesting and reduce pseudo replication. Statistical analysis was performed with JASP software (JASP Team, 2022) and is available at : https://osf.io/pt4br/overview, https://osf.io/uh73f/overview, https://osf.io/2wfy5/overview and https://osf.io/72jgz/overview.

### Exclusion criteria

Cells were excluded from AP firing or current analyses if a consistent increase or decrease in firing frequency or recorded current, respectively, was observed across pre-light sweeps. Individual sweeps were excluded from firing analysis when spontaneous activity between depolarizing steps obscured evoked firing, and from Vm analysis when spontaneous APs occurred during LED stimulation.

## Results

### Blue light produces a long-lasting reduction of the firing activity of mouse cortical neurons

To explore the impact of repetitive 5s blue LED stimulation at 19 mW on cortical neurons from adult male mice, we conducted current clamp recordings following the protocol outlined in Fig 1A. A negative current step was employed to monitor membrane resistance (Rm), followed by two depolarizing steps inducing firing activity. The first depolarizing step was associated with LED stimulation during the light period. Additionally, a test LED pulse was utilized to assess the acute modification of membrane potential by light (refer to Supplementary Fig 1 for additional details). As depicted in Fig 1B, the LED caused a slight increase in average tissue temperature of 1.8° ± 0.5 C (tested on separate slices), which returned to baseline within 20 s after the end of illumination (Supplementary Fig 4). Ten repetitions of LED stimulation resulted in a long-lasting decrease in firing activity (39% ± 27% of the pre-LED activity, p = 6 x 10^-4^; DF=16, N=17 neurons from 12 mice, Fig 1Ci). Similar effects were observed when six repetitions of LED stimulation were applied (Fig 1Cii, 37 ± 29 % of the pre-LED activity, p=0.015, DF=5, N=6 neurons from 6 mice). Extended recording time in these experiments highlighted the persistence of the firing decrease. An overall reduction to 38% ± 23% of the pre-LED activity was observed between 10 and 16 minutes after the initiation of the first LED stimulation (p = 5 x 10^-5^; DF=22, N = 23 neurons from 18 mice, see Fig. 1Ciii). However, no significant change in firing activity was observed when only a single light pulse was delivered (Fig. 1C iv, p = 0.52; df=5, N = 6 neurons from 3 mice). Suggesting that the repetitions of LED stimulation is required to produce the long-lasting effect. To assess the potential contribution of spontaneous drift to the firing changes observed after light stimulation, we performed a series of recordings in which, after a stable baseline activity had been established, neuronal firing was monitored for an additional 10 min prior to LED stimulation. As shown in Fig. 1C (vi–vii), no average modification were observed (138 ± 63% of baseline activity, p=0.26, df=8, N=9 neurons from 4 mice. p=7×10^-4^, df=30 when compared to the pot-LED firing of panel Ciii) . Importantly, subsequent LED stimulation still produce firing decrease (-40 ± 30% from the drift value, df=8, N=,9 p= 0.04).

**Figure 1.**
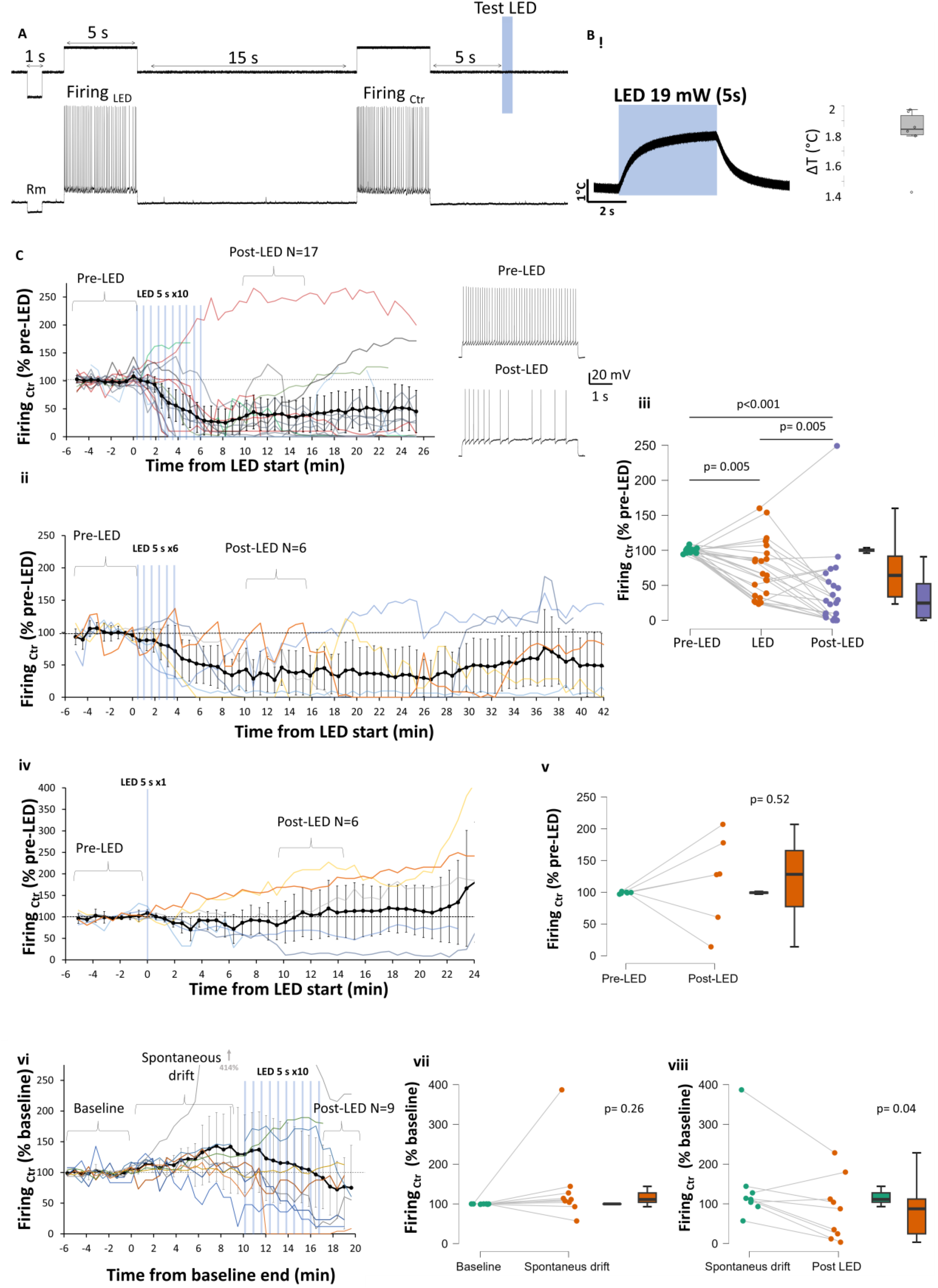
Light pulses induce a prolonged reduction in firing activity in mouse cortical neurons. (**A)** The current-clamp protocol used to study the effect of light on firing activity. This protocol included a negative current step to monitor membrane resistance, followed by two depolarization steps to elicit firing activity. The first depolarization (LED firing) occurred simultaneously with LED stimulation during the light period. In addition, a 1-second test LED pulse was used to evaluate the acute change in membrane potential caused by light. Top, injected current. Bottom, recorded voltage. (**B)** Transient increased tissue temperature following 5 seconds of LED stimulation. (**C)** Repeated LED pulses resulted in a long-lasting decrease in firing activity. **Ci** Time course of the modification of firing activity induced by 10 LED pulses. Representative traces of the light effect are shown on the right. **Cii**. Time plot illustrating the modification of firing activity induced by 6 LED pulses. **Ciii**. Statistical quantification of the light effect on firing activity in the three time periods. **Civ** Time plot illustrating the absence of modification of firing activity induced by 1 LED pulses. **Cv** Statistical quantification of the effect of 1 light pulse. **Cvi** Time plot illustrating the spontaneous evolution of firing activity after stable baseline period. **Cvii-Cvii** Statistical quantification of spontaneous evolution and the subsequent LED effect. (Post LED = 10-16 minutes); (ANOVA test followed by Holm post hoc comparisons). Black dots and bars represent the mean and 95% confidence interval, respectively. Colored lines depict the evolution of the firing rate across trials for each recorded neuron. Raw data and analysis are available at: https://osf.io/uh73f/

The light stimulation results in a significant and enduring average reduction of neuronal Rm (- 17 ± 15 MΩ, p=0.001, df=21,N=22 neurons from 18 mice, Fig 2A). Interestingly, the modification of firing produced by LED stimulation correlates with the Rm modification (r=0.52 [0.12;0.77], p=0.013, df= 19, N=21, see Fig 2Aiii). These results suggest that the LED effect on Rm accounts for approximately 27% of the variability in the LED effect on firing. On the other hand, the average resting membrane potential (Vrest) was not consistent affected by the light (Fig 2Bi and Bii) and the modification of this parameter only slightly correlate with the modification of firing (r=0.36, p=0.097, df=20, N=22, see Fig 2Biii). As previously reported ^9,10,12^, during test LED we observed a transient hyperpolarization of the membrane potential (Vm) (-0.31 ± 0.09 mV, p=1.4×10^-7^, df=37,, N=38 neurons from 22 mice, Fig. 2Ci), which reverts at -100 mV ± 8 mV (N=14 neurons from 9 mice, data not shown). Interestingly, this acute effect of light on Vm did not correlate with the long-term effect on firing (see Fig 2Ciii), suggesting that the two phenomena rely on different cellular mechanisms. Moreover, this lack of covariation also suggests that the observed variability in the light effect on neuronal firing is not due to methodological artifacts (e.g., the health of the recorded neuron or its depth in the slices).

**Figure 2.**
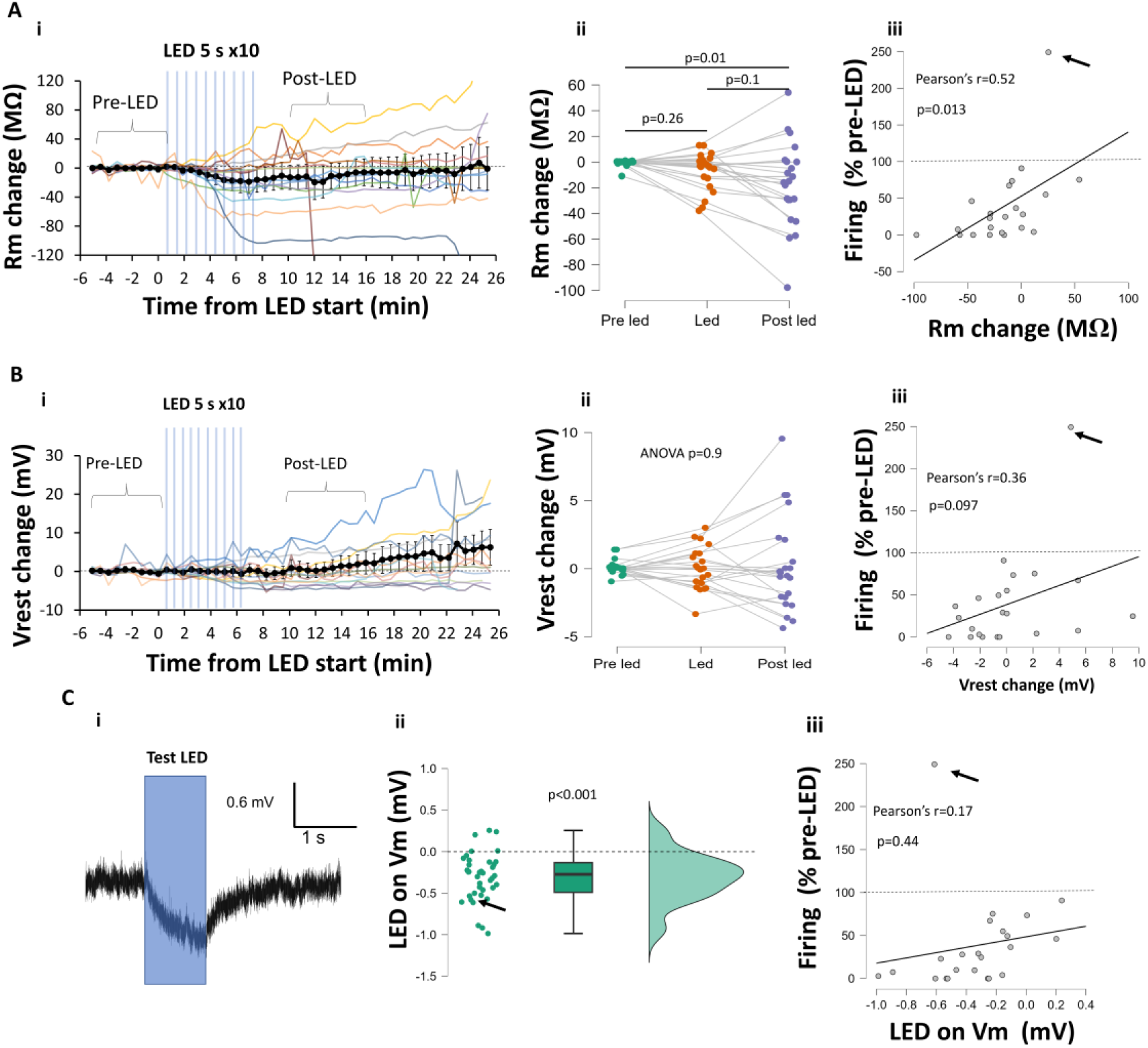
Changes in Rm explain part of the light effect on neuronal firing. **(A)** Repeated LED pulses resulted in a decrease in Rm. Ai. Time course of Rm modification induced by 10 LED pulses. Aii. Statistical quantification of the light effect on Rm in the three time periods. Aiii. LED-induced change in firing activity correlated with LED-induced change in Rm. **(B)** Repeated LED pulses on Vrest. Bi. Time course of Vrest modification. Bii. Statistical quantification of the light effect on Vrest in the three time periods. Biii. LED-induced modification of firing activity does not correlate with LED-induced modification of Vrest. **(C)** Light induces transient membrane hyperpolarization. Ci. Example of transient modification of Vm during light stimulation Cii. Quantification of the transient Vm change. Ciii. LED-induced modification of firing activity does not correlate with LED-induced transient modification of Vm. B and C: (post LED = 10-16 min) ANOVA test followed by Holm post hoc comparisons. Black dots and bars represent mean and 95% confidence interval, respectively. Colored lines represent single cell responses. Arrows depict the values for the neuron for wich light produced long lasting increase of firing. Raw data and analysis are available at : https://osf.io/uh73f/

To determine whether the reduction in neuronal firing was attributable to a light-induced increase in GABAergic transmission^16^ experiments were performed in the presence of 5 µM gabazine. As shown in figure 3A LED still produce a significant reduction of firing (44 ± 41% of baseline, N=6 neurons from 4 mice, p=0.02). Interestingly, this effect was associated with an increase in Rm (22 ± 15 MΩ, p = 0.02, Fig 3B), suggesting that the light-induced increase in GABAergic transmission may contribute to the decrease in membrane resistance observed in the absence of GBZ. The contribution of photodamage to the LED-induced effect was assessed by performing experiments in the presence of Trolox (100 µM), a water-soluble vitamin E analogue that scavenges several reactive radicals^17,18^ and has been reported to protect against light-induced phototoxicity in cell cultures and brain-slice preparations ^19,20^. As shown in Figure 3C, LED stimulation still significantly reduced neuronal firing in the presence of Trolox (27 ± 52% of baseline, N = 5 neurons from 2 mice, p = 0.02). In parallel, we assessed LED-induced free radical production by measuring superoxide anion production using an acetylated cytochrome c reduction assay. Superoxide anions was produced by the slices over the first 5 minutes following the transferring of the slices in the chamber (No-slices: 0.35 ± 0.10 pmol; With slices: 1.24 ± 0.26 pmol, p = 7 × 10⁻⁵ Fig. 3D). A small additional superoxide production was observed in the following 5 minutes only when exposure to LED was applied (LED: 0.29 ± 0.16 pmol, p = 8 × 10⁻⁴; CTR: −0.07 ± 0.13 pmol, p = 0.4, Fig. 3D). Yet the total amount of superoxide anion produced over the 10-minute period was not significantly different between the LED and Control group (p=0.9) No significant change was observed when LED stimulation was performed in ACSF alone (−0.07 ± 0.16 pmol, p = 0.4; data not shown). These findings suggest that cortical slices produce superoxide even in the absence of light, and that LED stimulation further increases its production.

**Figure 3.**
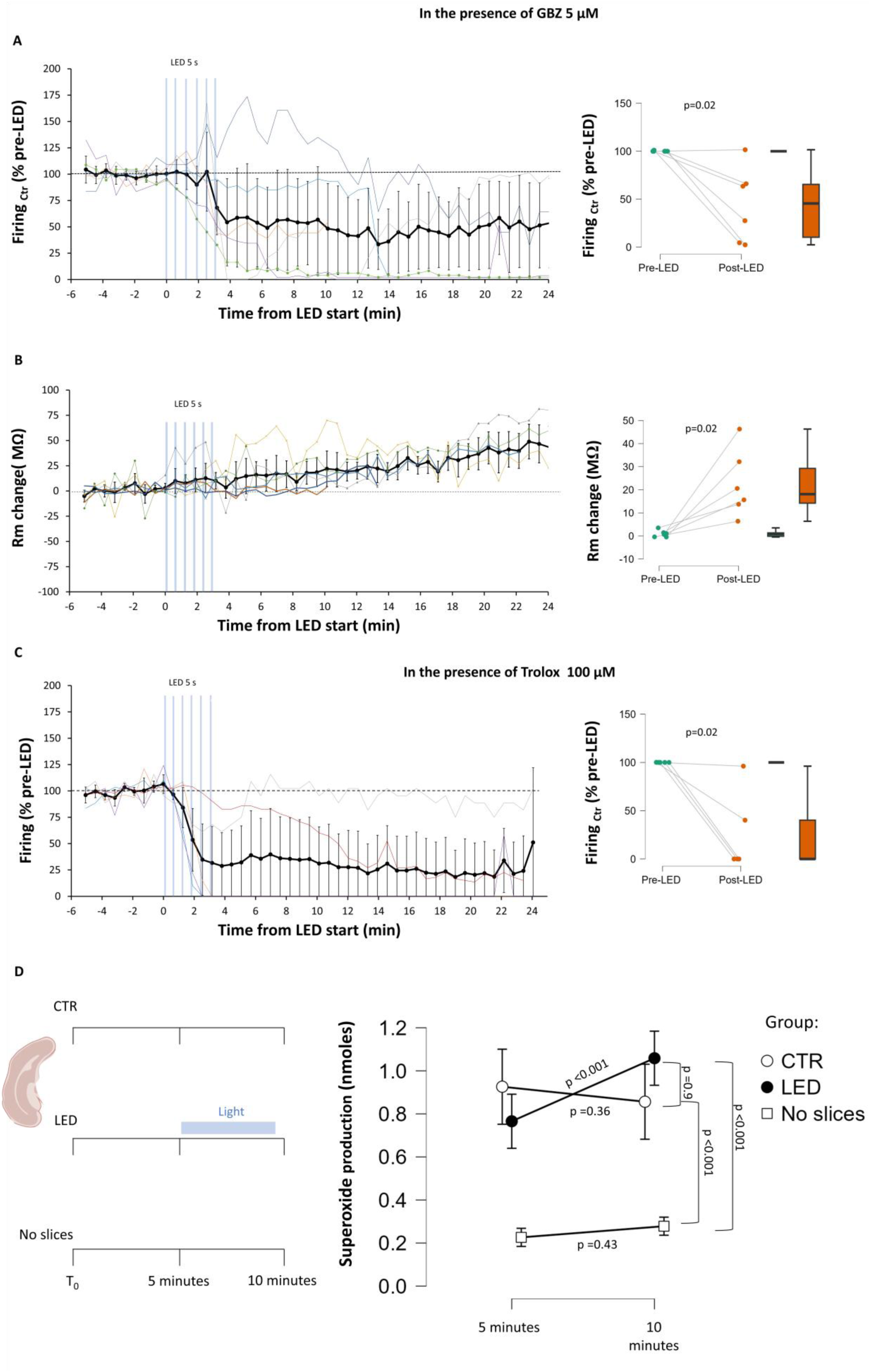
Role of GABAergic transmission and ROS generation in the effects of light stimulation on neuronal firing. **(A)** The reduction of firing activity produced by LED stimulation was not prevented by the presence of the GABA_A_ receptor antagonist Gabazine (GBZ). Left, Time plot illustrating the modification of firing activity induced by 6 LED pulses. Right, statistical quantification of the light effect on firing activity. **(B)** LED stimulation produce an increase of Rm in the presence of GBZ. Left, Time plot illustrating the modification of Rm by 6 LED pulses. Right, statistical quantification of the light effect on Rm. **(C)** The reduction of firing activity produced by LED stimulation was not prevented by the presence of the antioxidant Trolox. Left, Time plot illustrating the modification of firing activity induced by 6 LED pulses. Right, statistical quantification of the light effect on firing activity. **(D)** LED stimulation increases superoxide ion production in cortical slices. Left, schematic representation of the experimental protocol (see Supplementary Figure 3 for details). Right, quantification of superoxide ions production in the three experimental groups. p-values indicate post hoc comparisons between groups (independent-samples comparisons) and pairwise comparisons between time points within each group (time effect), with Holm correction for multiple comparisons. Colored lines in A, B and C depict the evolution of the firing rate and Rm across trials for each recorded neuron. Data analysis is available at https://osf.io/72jgz/overview

### Blue light induces lasting changes in voltage-dependent currents underlying APs

To investigate the involvement of voltage-gated sodium and potassium channels on the effect of light on firing activity, voltage-clamp recordings were performed to detect voltage-gated (VG) outward (putative K⁺) and inward (putative Na⁺) currents. These currents were elicited by experimentally modifying the membrane potential to mimic an AP waveform representative of cortical pyramidal neuron activity (Fig 4A, see Supplementary Fig 2 for more details). As shown in Fig 4B, LED stimulation produced a long-lasting reduction in VG-inward current amplitude (66% ± 28% of the pre-LED period, p=0.008, df=10, N=11 neurons from 5 mice). This effect developed slowly, requiring around seven minutes after the first LED stimulus to reach the maximum average reduction (Fig. 4Bii, Biii) and was associated with a slowing of VG-inward current kinetics (rise time from 4.4 ± 1.4 nA/ms to 3.1 ± 1.2 nA/ms; decay time from 3.4 ± 1.4 nA/ms to 2.2 ± 1.2 nA/ms, N=11 neurons from 5 male mice, Fig. 4Biv, 4Bv). Interestingly, the inward current was completely abolished in two recorded neurons (18%), which closely matches the proportion of neurons (17%) in which firing activity fully disappeared in current-clamp experiments. Although this close correspondence is noteworthy, the small sample sizes preclude determining whether the two proportions are related or whether their similarity is coincidental. It should be noted that in some neurons, the reduction of inward current following LED stimulation can revert back (see blue trace in Fig 4Bii). To isolate possible transient LED effects from the long-lasting LED effects, the VG-inward currents produced by the AP_LED_ were compared with those produced by the AP_Ctr_. As shown in Fig 4C (i-iii), the kinetics of the VG-inward current were slightly faster during LED stimulation than in the control condition (LED rise time = 4.5 ± 1.4 nA/ms, Ctr rise time = 4.2 ± 1.2 nA/ms; LED decay time = 3. 4 ± 1.4 nA/ms, Ctr decay time= 3.1 ± 1.4 nA/ms, N=11neurons from 5 male mice), while the VG-inward current amplitude was not altered (Fig. 4Civ).

**Figure 4.**
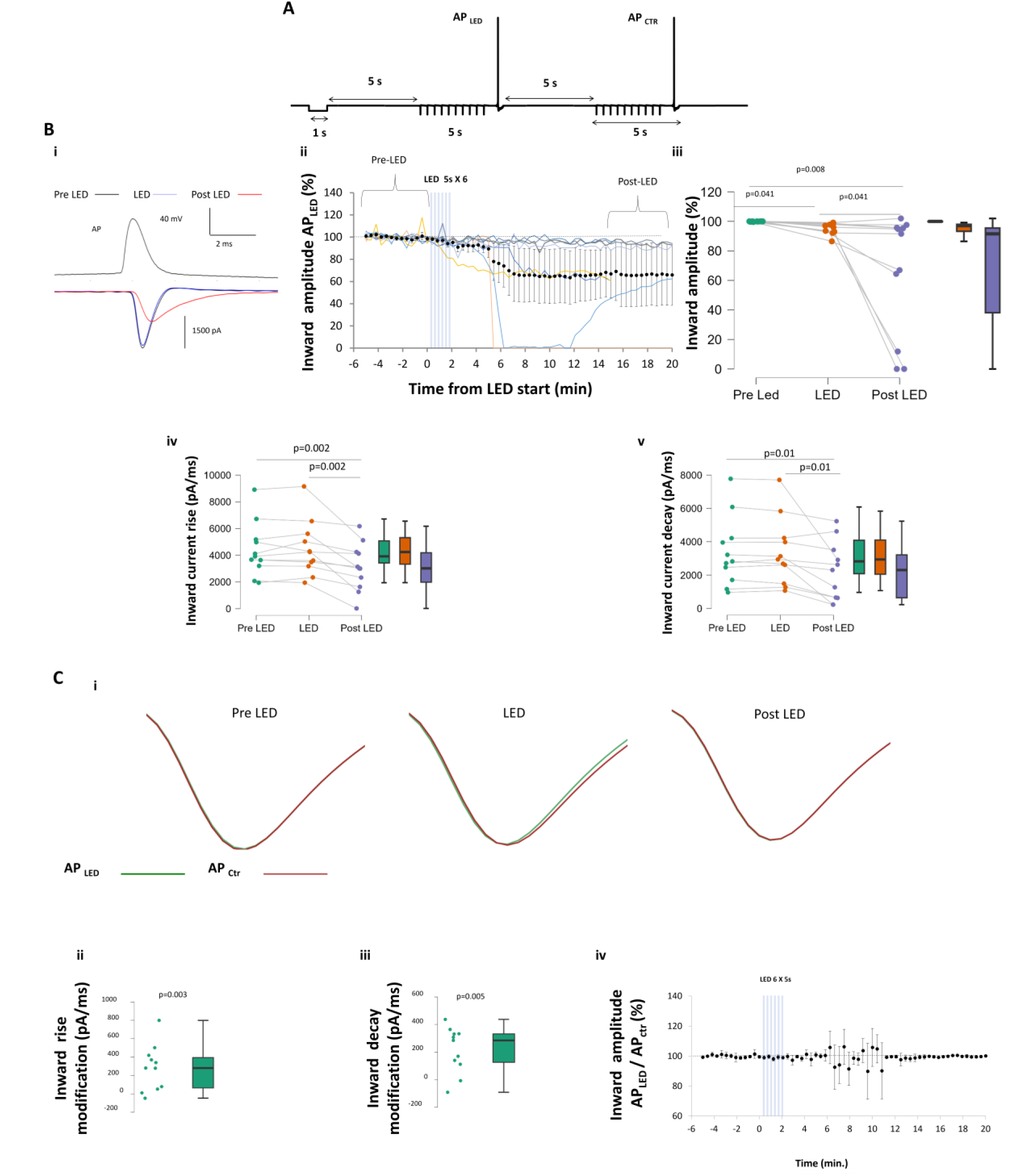
Light stimulation alters the voltage-gated inward current involved in AP generation. **(A)** Voltage clamp protocol used to isolate VG currents. Two consecutive AP potentials were generated by voltage command. AP_LED_ was generated simultaneously with light stimulation during the LED period. VG-currents were isolated from resistive and capacitive currents using the method shown in Supplementary Figure 2. (**B)** LED stimulation produces a sustained reduction in VG inward current amplitude and kinetics. Bi. Example of VG-inward current modification by LED stimulation. Bii. Temporal evolution of VG inward current amplitude. Black dots and bars represent mean and 95% CI, respectively. Coloured lines represent single-cell evolution. Biii. Statistical quantification of the average VG inward current amplitude in the three periods. Biv. Statistical quantification of the average VG inward current rise time in the three periods. Bv. Statistical quantification of the average VG inward current decay time in the three periods. (Post LED = 15-20 minutes); (Friedman test followed by Conover post hoc comparisons). (**C)** LED stimulation produces a short-term increase in VG-inward current kinetics. Ci. Example of overlap of VG-inward currents produced by AP_LED_ and APCtr in the three periods. Cii. Statistical quantification of the change in VG-inward current rise time during LED stimulation. Ciii. Statistical quantification of the change in VG-inward current decay time during LED stimulation. Civ. Temporal evolution of the ratio between AP_LED_ and APCtr VG-inward current amplitudes (n=11). Raw data and analysis are available at https://osf.io/pt4br/

As shown in Fig 5, LED stimulation induced a slight, long-lasting reduction in voltage-gated outward currents that did not reach statistical significance (average decrease of 45 ± 39 %, p=0.058 after Holm correction, df= 7, N=8 neurons from 4 male mice) (Fig 5Aii, Aiii). However the comparisons of AP_LED_ and AP_Ctr_ currents during LED period show a clear transient increase in the amplitude VG-outward current during the LED stimulus (131 ±7 % during LED,p= 4×10^-9^, df=7, N=8 neurons from 4 male mice, Fig 5B) showing a short-lasting effect of light on this parameter. In agreement with the current clamp experiment, the LED stimulation produced a long-lasting decrease in Rm, as monitored during voltage clamp recording (-60 ± 45 MΩ at 15’-20’, p=0.037, df=9, N=10 neurons from 6 male mice, Supplementary Figure 5 A,B,C). Importantly the effect of light on VG-inward current do not correlate with the effect of light on Rm (r=-0.22, p=0.59, df=7,N= 8 neurons from 5 mice; Supplementary Figure 5D); suggesting that the presence of two independent mechanisms (see discussion for theoretical implications). To assess the potential contribution of spontaneous drift to the changes observed after light stimulation, we performed a series of recordings in which, after a stable baseline activity had been established, inward current and Rm were monitored for an additional 18 min prior to LED stimulation. As shown in Supplementary Figure 6, no spontaneous drift was observed either in inward current (95 ± 10% of baseline activity, p = 0.13, df = 11, N = 12 neurons from 7 mice) or in Rm (7 ± 30 MΩ from baseline, p = 0.86, df = 8, N = 9 neurons from 7 mice). Importantly, at this 18-min time point, both the inward current and Rm modifications were significantly bigger in the stability-control experiments than in the experiments in which LED stimulation was delivered at time 0 (p=0.035, df=21 for inward current and p=0.04, df=16 for Rm). Subsequent LED stimulation still produced a significant decrease in inward current (supplementary figure 6 A iii) and Rm (supplementary figure 6 B iii) Taken together, these results suggest that light causes both transient and long-lasting changes in the biophysical properties of voltage-gated channels involved in the AP, as well as in channels involved in resting membrane permeability. Transient effects are likely to be produced by the light-induced temperature rise, leading to a faster activation and inactivation rate of the VG-inward current and an increase in the amplitude of the VG-outward currents^21^. Long-term changes include a decrease in the amplitude and kinetics of VG-inward currents, a heterogeneous effect on VG-outward current amplitude dominated by a decrease of the latter, and a predominant decrease in Rm.

**Figure 5.**
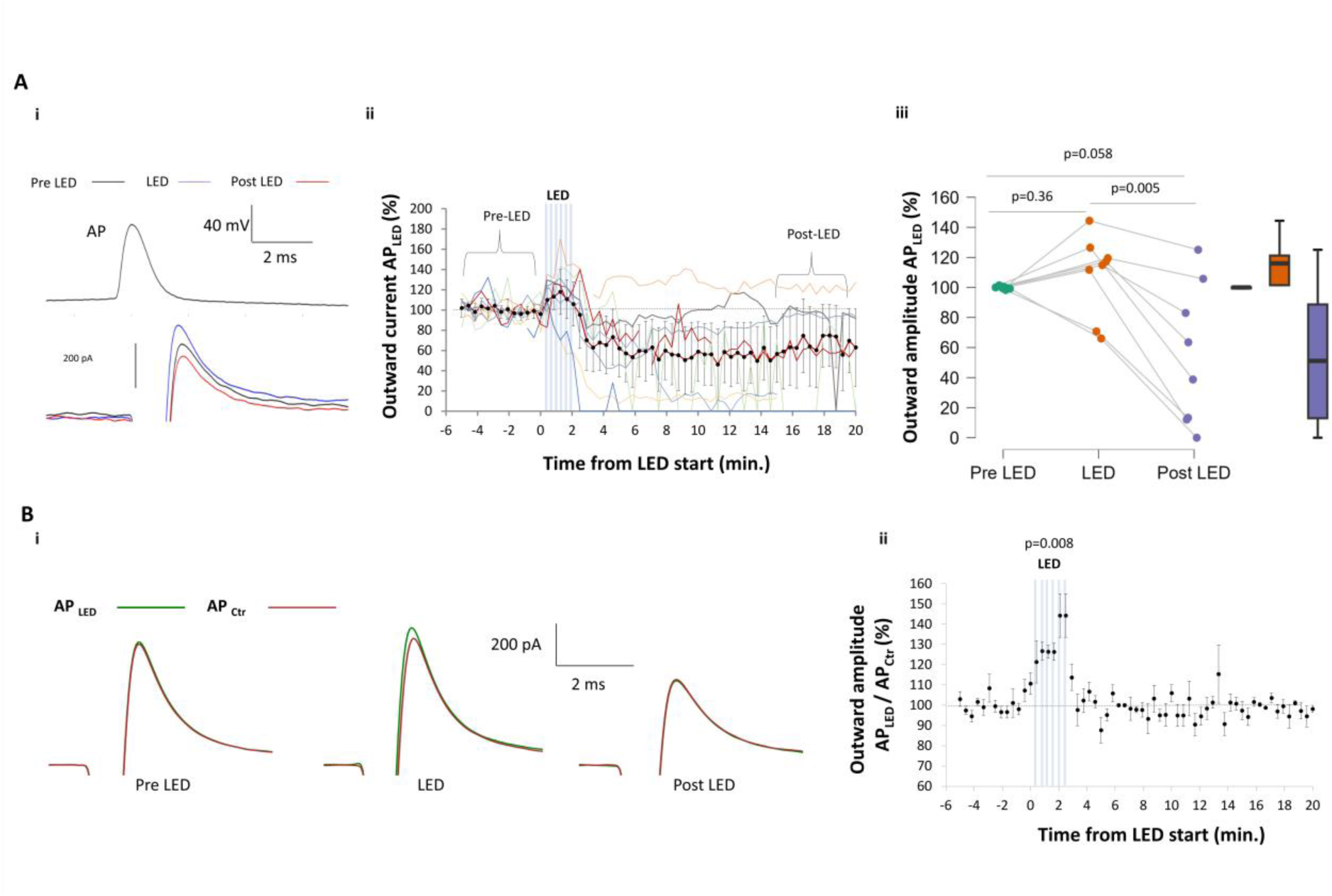
Light stimulation modifies the voltage-gated outward current involved in AP generation. (**A)** LED stimulation produces a long-lasting change in the amplitude of the voltage-gated outward current. Ai. Example of outward current modification by LED stimulation. Aii. Temporal evolution of VG outward current amplitude. Black dots and bars represent mean and 95% CI, respectively. Colored lines represent single-cell evolution. Aiii. Statistical quantification of the average outward current amplitude in the three periods (Friedman test followed by Conover post hoc comparisons). (**B)** LED stimulation produces a transient increase in VG outward current amplitude. **Bi.** Example of overlap of VG outward currents produced by AP_LED_ and APCtr in the three periods. **Bii.** Temporal evolution of the ratio between AP_LED_ and APCtr VG outward current amplitudes. Raw data and analysis are available at https://osf.io/pt4br/

The theoretically expected effects of changing these three parameters on the direction of firing change are illustrated in Fig 6A. In order to evaluate their reciprocal contribution to the light-induced long-lasting decrease of firing, we plotted the modification of evoked firing activity, that was monitored before and at the end of voltage clamp protocol, as a function of the modifications of the VG-currents and the Rm (Fig 6B). This analysis suggests that the light-induced decrease of firing rate is a multifactorial process with the modification of VG-inward current being the principal actor. A large decrease in VG inward currents was always associated with a large decrease in firing rate. On the other hand, a small decrease in inward currents was associated with either a strong, medium, small or no decrease in firing rate, depending on the direction and magnitude of the modification of VG-outward currents and Rm.

**Figure 6.**
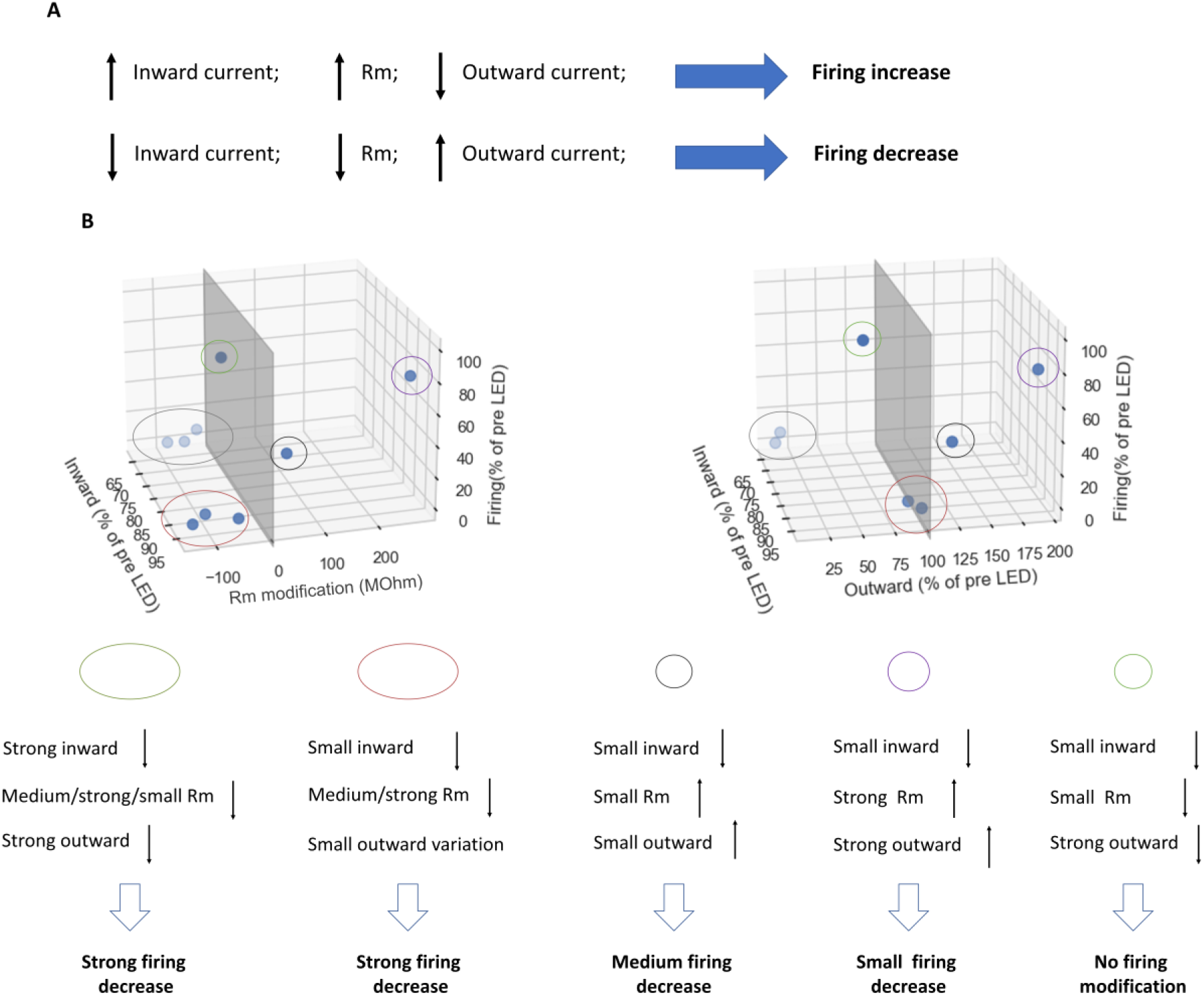
Relative contribution of the light-induced changes in VG-currents and Rm to the modification of neuronal firing rate. (**A)** Expected impact of VG-currents and Rm modifications on the direction of firing rate change. (**B**) Top, 3D plot illustrating the light-induced modification of firing rate as a function of the light-induced modification of VG-inward, VG-outward and Rm. Bottom, Classification of the effect on firing activity associated with the different combinations of observed changes in VG currents and Rm.

### Heterogenous effect of blue light stimulation on cortical human neurons

The effect of blue light stimulation on human cortical neurons was investigated using a current clamp configuration, following the same protocol as for mouse neurons (Fig 1A). Tissue samples came from nine patients differing in age, sex, pathology, and the explanted cortical region. LED stimulation resulted in a heterogeneous modification of firing activity in the fifteen neurons recorded. Specifically, there was a significant long-lasting reduction of firing in 8 neurons (55%) and a significant long-lasting increase in firing in 4 neurons (27%) while three neurons were unaffected (20%) (Fig 7A). Overall, the light-induced effect was significantly smaller in human than in mouse neurons when neurons were treated as independent observations (−12 ± 43% of pre-LED firing in human neurons; df = 50, p = 0.025). This difference remained significant when the effects were aggregated across neurons from the same subject (df= 26, p=0.023). The proportions of excited and inhibited neurons also differed significantly between species when neurons were treated as independent observations (χ² test, χ² = 31, df = 2, p = 1 × 10⁻⁷, N = 35). However, this difference was no longer significant when the nested structure of the data was taken into account (GLM, df = 1, p = 0.35). To rule out the possibility that this difference resulted from the longer interval between tissue removal and slice preparation in human samples, we performed additional experiments in which mouse brains were kept in cold, oxygenated solution for 20 minutes before slicing, thereby reproducing the delay associated with human tissue preparation. As shown in supplementary figure 7 this manipulation did not alter the light induced reduction of firing previously observed in mouse neurons (28 ± 31 % of the pre-LED activity, p=0.01, df=4, N=5 neurons from 2 male mice) . Moreover exploratory analysis did not reveal any distinguishable biophysical properties between mice and human neurons explaining this difference (for more details, see https://osf.io/2wfy5/files/45kp3). Moreover, exploratory analysis showed that the proportion of excited neurons was significantly higher in tissue from female patients compared to male patients (83% vs 12%, respectively, p=0.008, df=1, χ² test, Fig 7B, see also Supplementary Fig 9A-B). This difference remained significant after accounting for the nested structure of the data (generalized linear mixed model, χ² = 5.3, df = 1, p = 0.022). suggesting that sexual dimorphism could be the cause of the heterogeneity in the effect of LED stimulation on firing in the human sample. Though, ultimately, due to the exploratory nature of this analysis this conclusion is speculative and would require more dedicated investigation in subsequent studies. Other factors, including patient age and underlying pathology, may also act as confounding variables that could explain the sex-related differences in the effects of light on neuronal firing. However, exploratory analyses revealed only a moderate correlation between age and the light-induced effect and no significant effect of donor pathology (Fig. 7C–D). Furthermore, neither other neuronal biophysical properties nor the magnitude of the light-induced change in Vm correlated with patient age (Supplementary Fig. 8). Finally, LED stimulation did not induce any detectable changes in Rm and Vrest in human neurons (Supplementary Fig. 10)To test whether the sex difference in light effects is also present in mice, ten additional cortical neurons were recorded from seven female mice. As shown in Supplementary Figure 9 C-D, light stimulation produces a similar long-lasting inhibition of firing activity in mouse neurons from both sexes (sex effect p = 0.87; BF01 = 3.5).

**Figure 7.**
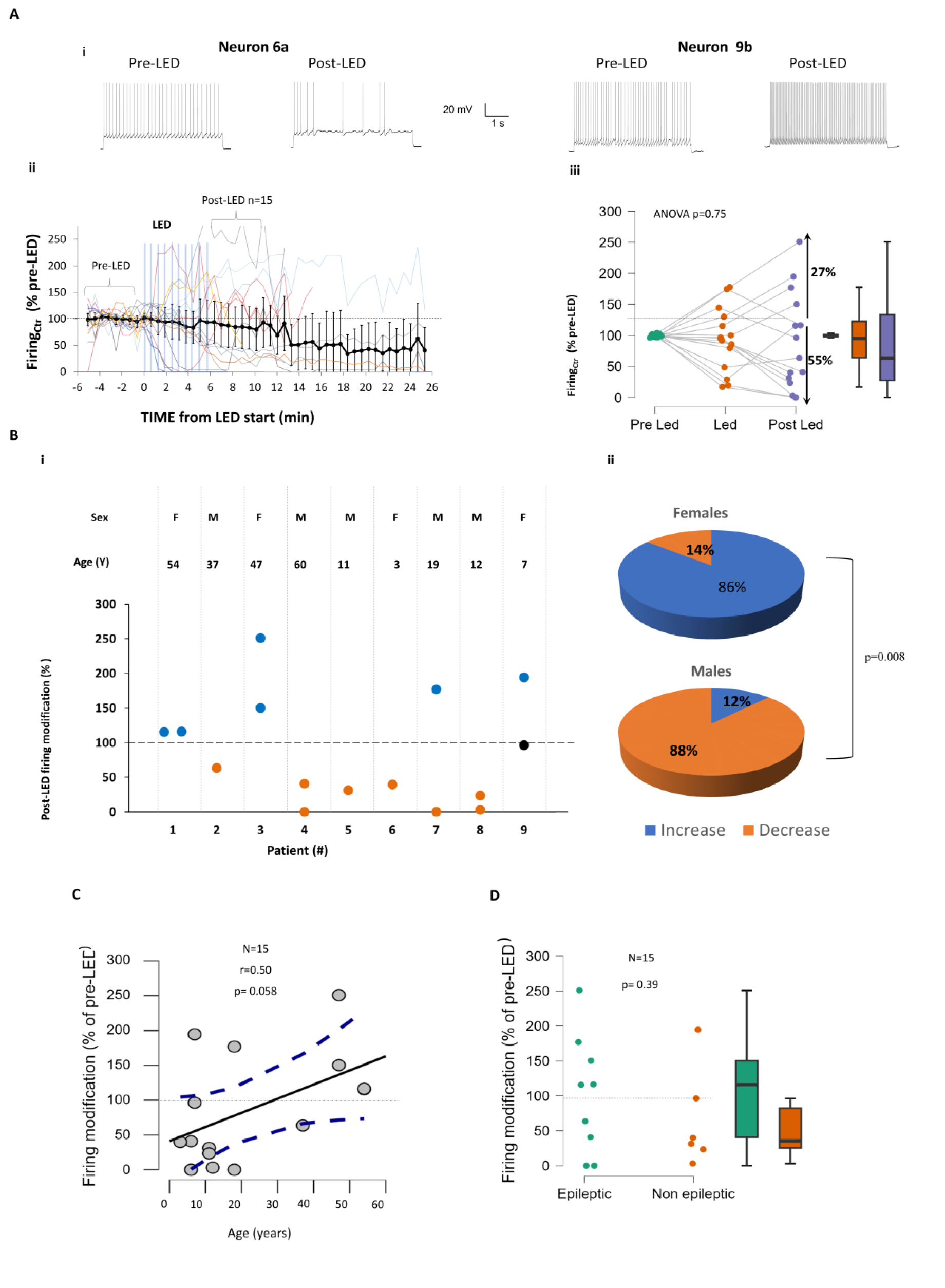
Light Pulses Induce a Prolonged and Heterogeneous Modification in Firing Activity in Human Cortical Neurons. **(A)** Repeated LED pulses induce a long-lasting decrease in firing activity in 55% of recorded neurons and an increase in firing in 27% of recorded neurons. Ai Representative traces showing the effect of light on two different neurons. Aii Time course of the firing activity modification induced by 10 LED pulses. Aiii Statistical quantification of the light effect on firing activity across three time periods (Post LED = 6-11 minutes). **(B)** Sexual dimorphism in the long-lasting effect of light on the firing modification of human neurons. Bi Long-lasting LED-induced firing modification in neurons from different patients. Bii Long-lasting LED effect on firing activity as a function of the sex of the donor patient. **(C)** Linear correlation between the effect of light and patient age **(D)** Long lasting effect of light on neuronal firing do not depend on patient pathology .Blue, orange, and black dots in B respectively represent neurons that showed a significant increase, a significant decrease, or no change in firing activity. Data are presented as mean ± 95% CI. Raw data and analysis are available at: https://osf.io/2wfy5/.

Interestingly, human neurons also differ from mouse neurons in their acute response to light on membrane potential. In 4 out of 23 (17%) human neurons, blue light produced a depolarization of the membrane potential (Fig 8Ai, Aii), an effect that was never observed in mouse neurons in the present study (Fig 8B, p<0.001, χ² test) nor previously reported in other studies. The reversal potential of this effect was estimated at -35 mV (N=1). When these neurons were excluded from the analysis, a global membrane hyperpolarization was observed (Vm = -0.22 mV ± 0.094 mV, p < 0.001; Fig 8Aiii), consistent with the finding in mouse neurons (Bayesian statistic, Mice vs. Human, BF01 = 1.9). The proportion of neurons exhibiting the depolarizing effect was slightly larger in females compared to males, although this difference did not reach significance (25% in females vs. 9% in males, p=0.31 χ² test, BF01=1.76)

**Figure 8.**
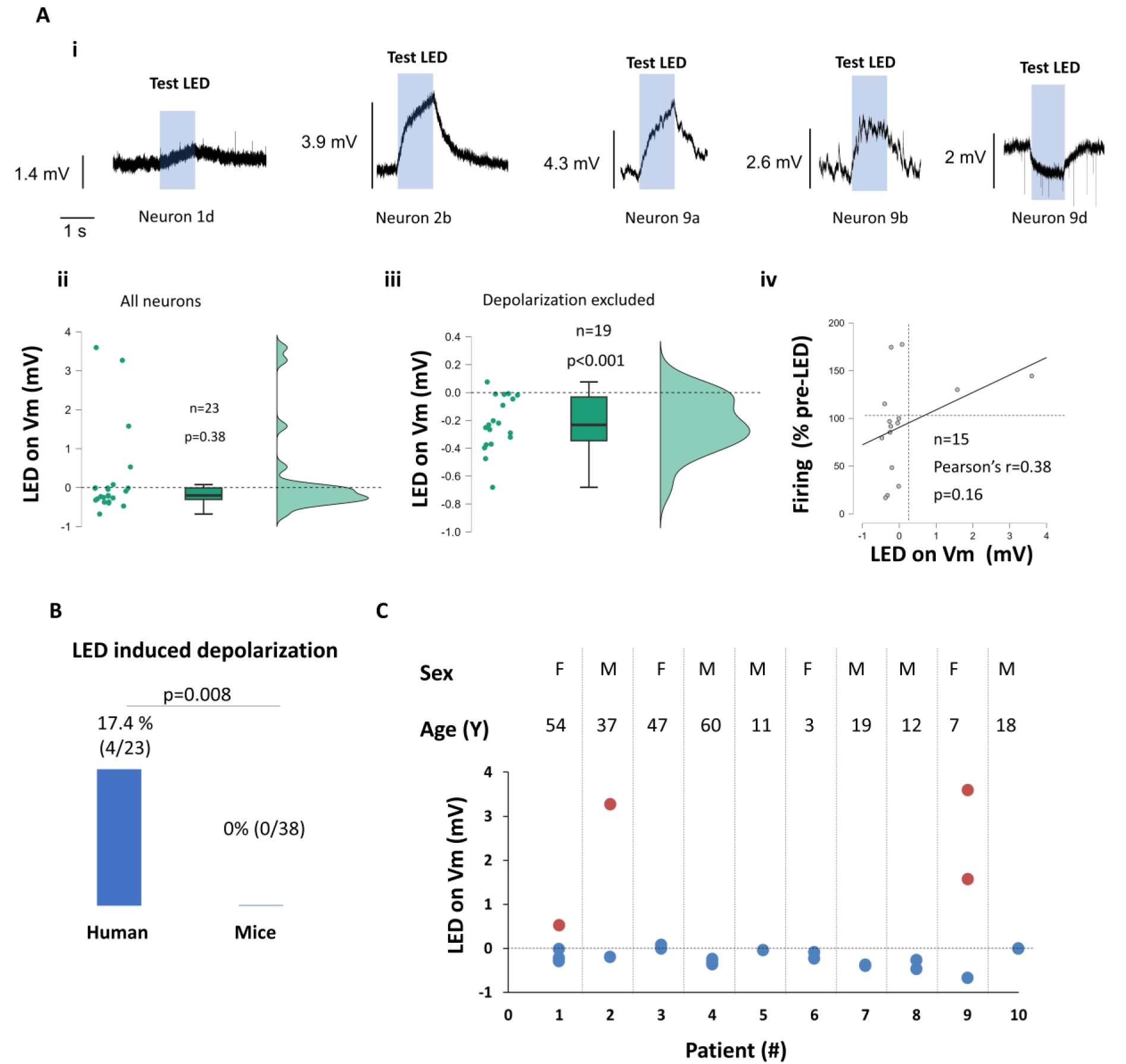
Heterogeneous modification of the membrane potential by LED stimulation in human neurons. **(A)** Light-induced membrane potential modification in human neurons. **Ai** Raincloud plot showing the effect of light on the entire sample. **Aii** Average traces illustrating the light-induced membrane potential modification in four neurons showing depolarization and one neuron showing hyperpolarization (average effect produced by ten stimulations). **Aiii** Raincloud plot of the light effect excluding the four depolarizing neurons. **(B)** The proportion of light-induced membrane depolarization was significantly higher in neurons from humans compared to those from mice. **(C)** LED-induced Vm modification in neurons from different patients (in red the four neurons showing Vm depolarization). Raw data and analysis are available at : https://osf.io/2wfy5/

## Discussion

This study investigated whether light stimulation induces sustained changes in neuronal excitability. Repeated blue light exposure reduced evoked firing by approximately 60% in cortical neurons from male and female mice, with effects persisting for tens of minutes after stimulation. Human cortical neurons showed more variable responses, with increased excitability in a subset, particularly those from female subjects. These prolonged effects may involve changes in passive membrane properties, including Rm and active conductances mediated by voltage-gated Na⁺ and K⁺ channels.

### Light Pulse Parameters Driving Sustained Suppression of Neuronal Activity

Previous studies reported reduced firing in several CNS neurons during visible-light stimulation, but activity generally recovered rapidly after stimulation ceased. This was particularly evident in Owen et al.^10^, who used 15 mW green light. In contrast, our study found persistent firing inhibition following 19 mW blue-light stimulation. Differences in wavelength and power may contribute to this discrepancy, although other methodological differences cannot be excluded. While a single pulse was insufficient to induce persistent changes, six and ten light pulses produced similar long-lasting inhibition, suggesting that relatively few pulses may suffice to achieve maximal firing inhibition. Firing continued to decline after the sixth pulse (Fig. 1Cii), suggesting that light triggers an effect that develops over several minutes.

### Implications of Phototoxicity and ROS for the Long-Lasting Effects of Light

Although the energy carried by visible light is insufficient to cause ionization and directly damage DNA or other cellular structures, it can still trigger photochemical reactions and oxidative stress, leading to cellular damage. For instance, blue light can induce the production of reactive oxygen species (ROS) in culture media, which has been associated with increased gene expression and neuronal death^22–24^. However, our findings do not support a marked acute cytotoxic effect. Although light stimulation altered several biophysical properties of recorded neurons, there was no clear evidence of compromised cell viability in the short term. Neuronal death is typically associated with rapid membrane depolarization; however, only modest depolarization was observed following light exposure. In addition, the light-induced change in membrane resistance (−9%) was comparable in magnitude to variations reported under other experimental conditions^25,26^ Furthermore, although firing activity was reduced, it remained detectable in most neurons and showed partial or complete recovery over time in some cases. ROS are not exclusively associated with oxidative stress and cellular damage. At controlled levels, they also act as signaling molecules involved in neuronal function^27–29^,including modification of neuronal excitability^30–32^ Here, we show that LED stimulation increased superoxide anion production by only 38% relative to the initial production measured in the presence of cortical slices, consistent with the basal superoxide production reported before^33^. Trolox, an antioxidant capable of scavenging several ROS, did not prevent the LED-induced decrease in neuronal firing. Although these data argues against a major contribution of ROS in the firing reduction, they do not completely exclude the possibility that superoxide anions contribute to the induction of the LED effect. Further investigations are therefore required to clarify the role of ROS in the long-lasting effects of light on neuronal excitability.

### Cellular mechanisms of light effect on firing activity

Among the mechanisms potentially underlying the persistent reduction in neuronal firing, we highlight decreased membrane resistance and reduced inward currents involved in action potential depolarization. The decrease in membrane resistance appears to depend on GABA_A_ receptors and may reflect enhanced tonic GABAergic conductance potentially linked to increased GABA_A_ receptor responsiveness following light stimulation ^16^. The increase in membrane resistance observed in gabazine suggests an additional, GABA_A_-independent mechanism that remains to be identified. The persistent reduction in inward current amplitude, accompanied by increased rise and decay times, suggests altered Na⁺ channel permeability or gating properties, potentially affecting action potential generation in cortical neurons^34^. In contrast, inward currents became faster during LED stimulation, while the amplitude of the outward, putative K⁺ current increased. These acute effects may result from transient warming during illumination^21^. Although temperature-induced changes in neuronal activity have been reported to reverse rapidly after warming ceases^10,21^, a contribution of repeated warming to the persistent effects observed here cannot be excluded. The measured temperature increase (1.8°C) was within the range of physiological temperature fluctuations (2–3°C)^35^, however, this does not rule out temperature-related effects on neuronal function, and further studies are needed to clarify its contribution.

### Species and sex differences in light effects

We identified two main differences in light responses between mouse and human cortical neurons. First, a long-lasting increase in firing activity was observed in approximately 30% of human neurons, whereas this effect was largely absent in mice. This apparent species difference may be influenced by a sex-specific effect observed only in human tissue, with neurons from female patients more frequently exhibiting increased excitability in response to light than those from male patients. However, it is important to note that these sex-based difference are both tentative and speculative due to the limited data pool. Second, an acute depolarizing response to light was detected in a subset of human neurons (17%) but was not observed in mice. These differences may reflect known divergences in the biophysical properties of human versus rodent neurons^36–38^. While previous studies have suggested that light-induced hyperpolarization primarily involves increased permeability of inwardly rectifying K⁺ (Kir) channels^10^, the depolarizing responses observed in some human neurons are more consistent with modulation of channels permeable to Na⁺ and/or Ca²⁺, such as Na⁺ leak channels or transient receptor potential (TRP) channels.

### Limitations and caveats

A key limitation concerns the interpretation of apparent sex-dependent differences in human neuronal responses to light. Although statistically supported, this finding arises from an exploratory analysis in which multiple factors, including age, pathology, and intrinsic electrophysiological properties, were examined as potential sources of variability. Differences in age and cortical sampling between male and female subjects may also confound the results, limiting the strength of this conclusion. Replication in larger, more controlled cohorts is therefore requiredAnother caveat concerns the estimation of voltage-gated currents associated with AP generation. Because voltage-gated Na⁺ channels are located predominantly at extrasomatic sites, particularly in the axon initial segment, somatic voltage-clamp recordings are subject to space-clamp limitations that may affect current estimates. This is particularly relevant here because LED stimulation reduced somatic membrane resistance, potentially modifying electrotonic coupling between the soma and axon initial segment and thereby altering the voltage experienced by Na⁺ channels during somatic voltage clamp. However, LED-induced changes in inward current did not correlate significantly with changes in membrane resistance (Supplementary figure 5D), arguing against the latter being the primary determinant of the observed decrease in inward current. Nevertheless, other passive and morphological properties, including axial resistance, membrane capacitance, axonal geometry, and the spatial relationship between the soma and axon initial segment, could also influence space-clamp conditions and contribute to the observed effect^39^. Direct demonstration of a light-induced effect on Na⁺ channel function would require complementary experiments in a simplified, electrically compact system, such as heterologous expression, although this would not fully reproduce the native neuronal environment. The voltage-clamp results could also be affected by LED-induced changes in the voltage dependence of passive currents, potentially overestimating passive currents and underestimating voltage-gated currents. However, this seems unlikely, as it would require LED stimulation to alter passive-current voltage dependence rather than simply its amplitude. Technical constraints inherent to patch-clamp recordings, including cytoplasmic dialysis and seal instability ^26,40^, may introduce variability and affect the accurate quantification of light-induced effects, particularly over prolonged recordings. Alternative approaches will therefore be needed to assess more reliably the persistence of the firing changes described here (see following section). Finally, limited visible-light penetration into biological tissue remains a major challenge for its potential therapeutic application in the human brain.

### Perspectives and conclusions

This study demonstrates that visible light can induce sustained changes in neuronal activity, providing a basis for investigating its therapeutic potential. Several directions warrant further investigation.

First, the effects of light on human neurons should be confirmed in larger, standardized cohorts, particularly regarding the apparent sex-related differences and potential influence of pharmacological treatments. Human brain organoids^41^ could improve tissue availability, reduce variability, and facilitate long-term efficacy and safety assessments.

Second, whether the sustained reduction in neuronal excitability observed in brain slices can be reproduced in vivo remains to be established. Although transient inhibitory effects of light have been reported in animal models^10,11^, electrophysiological recordings and behavioural assessments are needed to determine their persistence and functional relevance.

Third, the duration and reversibility of light-induced modulation should be characterized, including whether inhibition can be maintained or re-induced following recovery. Neuronal cultures, organotypic slices, organoids, and in vivo preparations with chronic recordings could help address these questions and assess long-term safety.

Fourth, sustained inhibition may be relevant to neurological disorders involving neuronal hyperexcitability, including epilepsy, migraine^42,43^, autism spectrum disorder^44,45^, and potentially schizophrenia^46^. Given the established use of optogenetic strategies to suppress pathological activity in epilepsy models^47–53^, it would be particularly interesting to determine whether light alone, without exogenous opsin expression, can exert antiepileptic effects.

Finally, stimulation parameters require optimization. Beyond the 5-s blue-light pulses used here (19 mW), future studies should investigate different wavelengths, intensities, and durations. Longer wavelengths may improve tissue penetration and enable modulation of deeper brain regions^54^.

Overall, these findings support further investigation of visible light as a strategy for modulating neuronal excitability. Studies in human systems and in vivo models will be essential to establish its efficacy, safety, and therapeutic potential.

## Abbreviations

ACSF: artificial cerebrospinal fluid
AP: action potential
LED: Light-Emitting Diode
GBZ: gabazine
Rm: membrane resistance
R_A_: access resistance
ROS: Reactive Oxygen Species
VG: voltage-gated
Vm: membrane potential
Vrest: average resting membrane potential.

## Data availability

Data are available via the Open Science Framework website (https://osf.io/6jqye/)

## Acknowledgements

We thank Belkacem MESSAOUDI for technical support. ChatGPT (OpenAI) was used for language editing and improving the clarity of the manuscript. All modifications were reviewed and approved by the authors.

## Supplementary figures

**Supplementary Figure 1:**
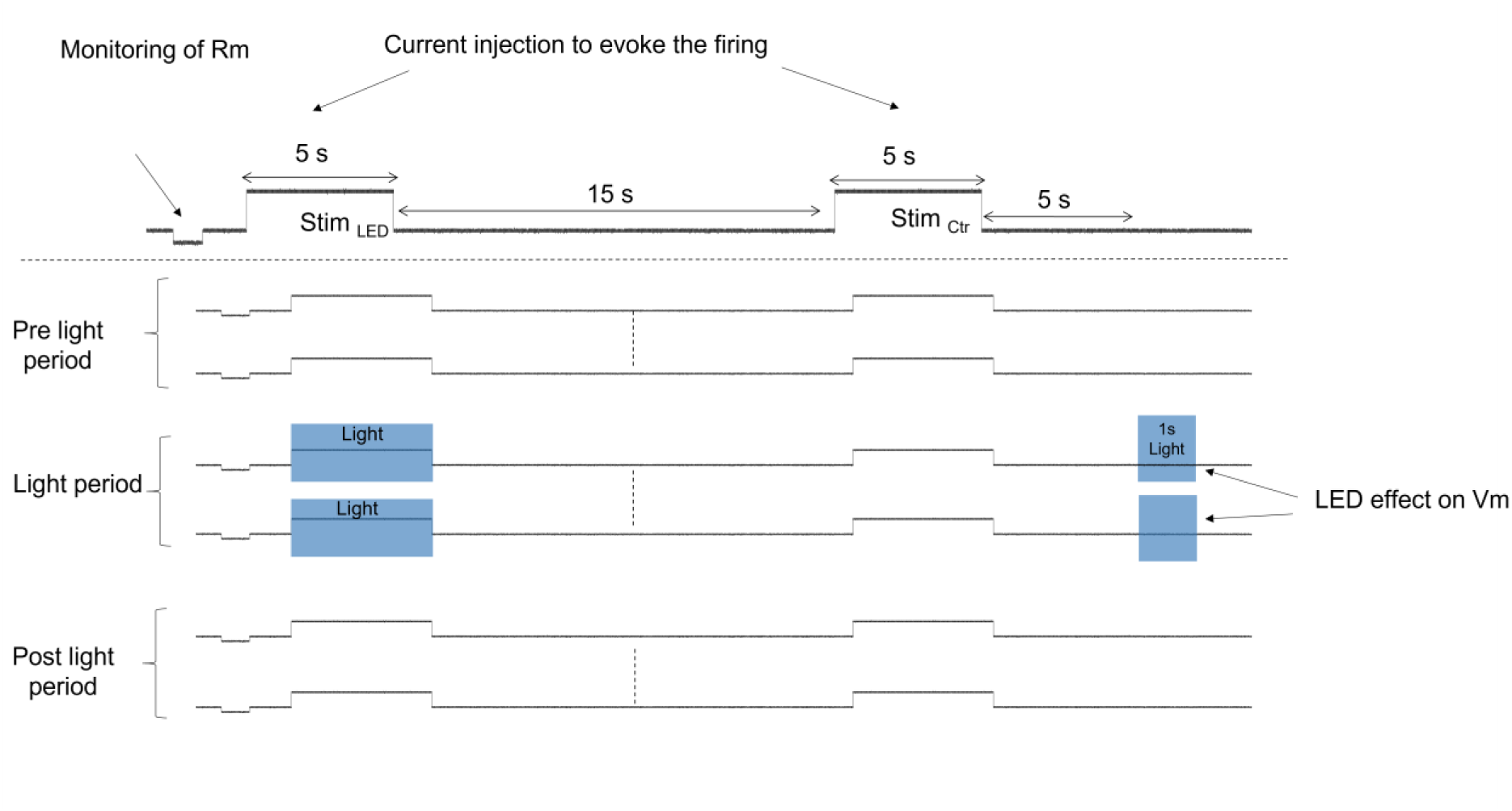
Current-clamp protocol to assess the light-induced long-lasting modifications of firing activity and membrane properties of cortical neurons. Top: timing of current injection during sweep acquisition. Bottom: sequence of protocol repetition (9 sweeps during the pre-light period, 10 or 6 sweeps during the light period). LED stimulation is depicted by the blue boxes.

**Supplementary Figure 2.**
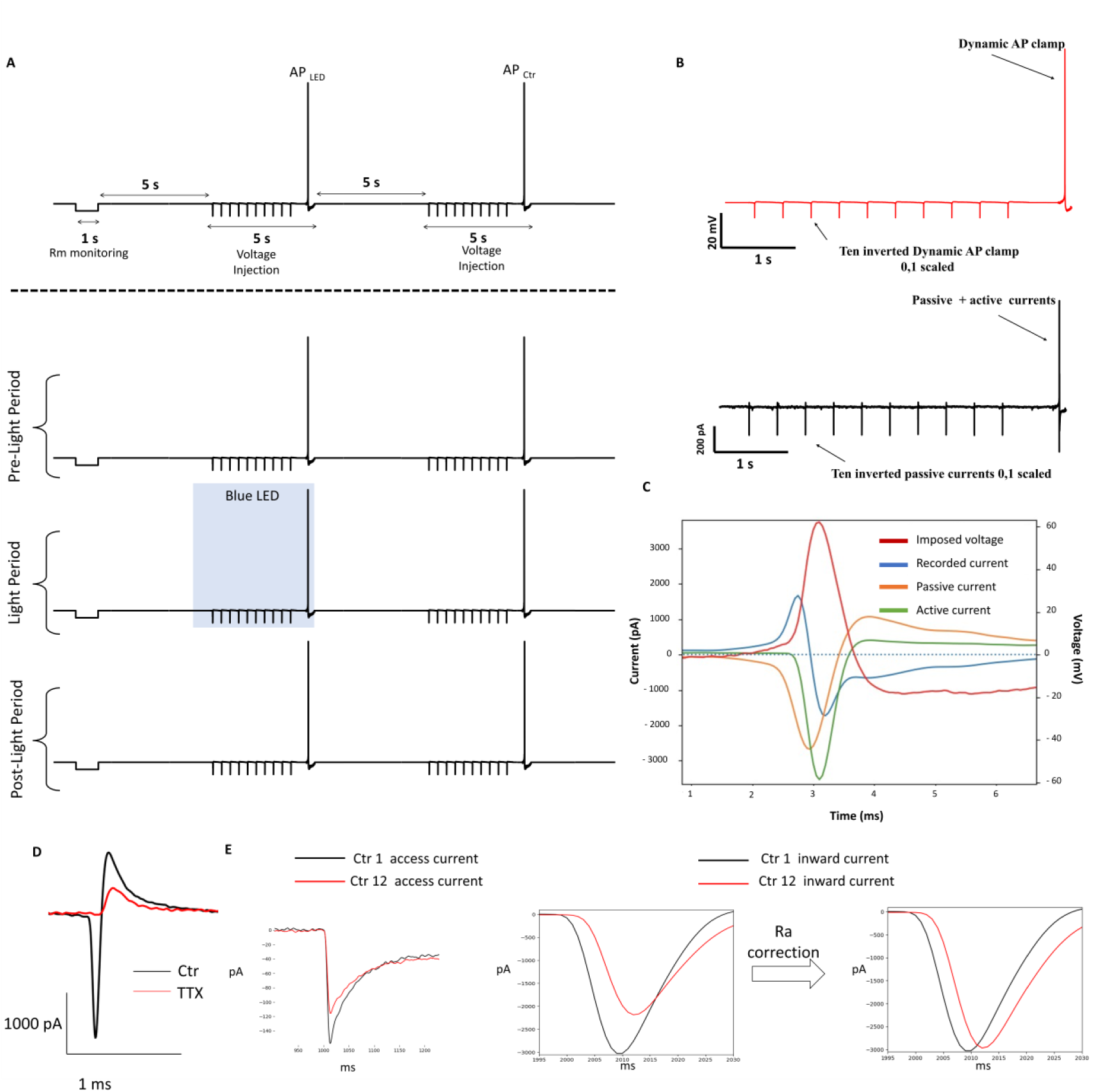
Voltage-clamp protocol to assess the light-induced long-lasting modifications of voltage-dependent currents involved in the action potential generation of cortical neurons. A) Top: Timing of voltage modification during sweep acquisition. Bottom: Sequence of protocol repetition (13 sweeps during the pre-light period, 6 sweeps during the light period). LED stimulation is depicted by the blue boxes. B) Top: Voltage modification applied to the recorded neurons. Bottom: Recorded currents. C) Example of the isolation of active currents during fake action potential generation (in red). Total passive currents (sum of the ten inverted passive current, in orange) were removed from the recorded currents (in blue), leaving the voltage-dependent currents (in green). D) The inward current was abolished by tetrodotoxin (TTX) 1 µM. E) Example of the effect produced by correcting the current trace to account for changes in access resistance. Left: reduction of the access current between the first and the twelfth sweep recorded under control conditions; center: impact of this reduction on the calculated inward current; right: effect of the Ra correction on the inward currents. Holding voltage =-70 mV.

**Supplementary Figure 3.**
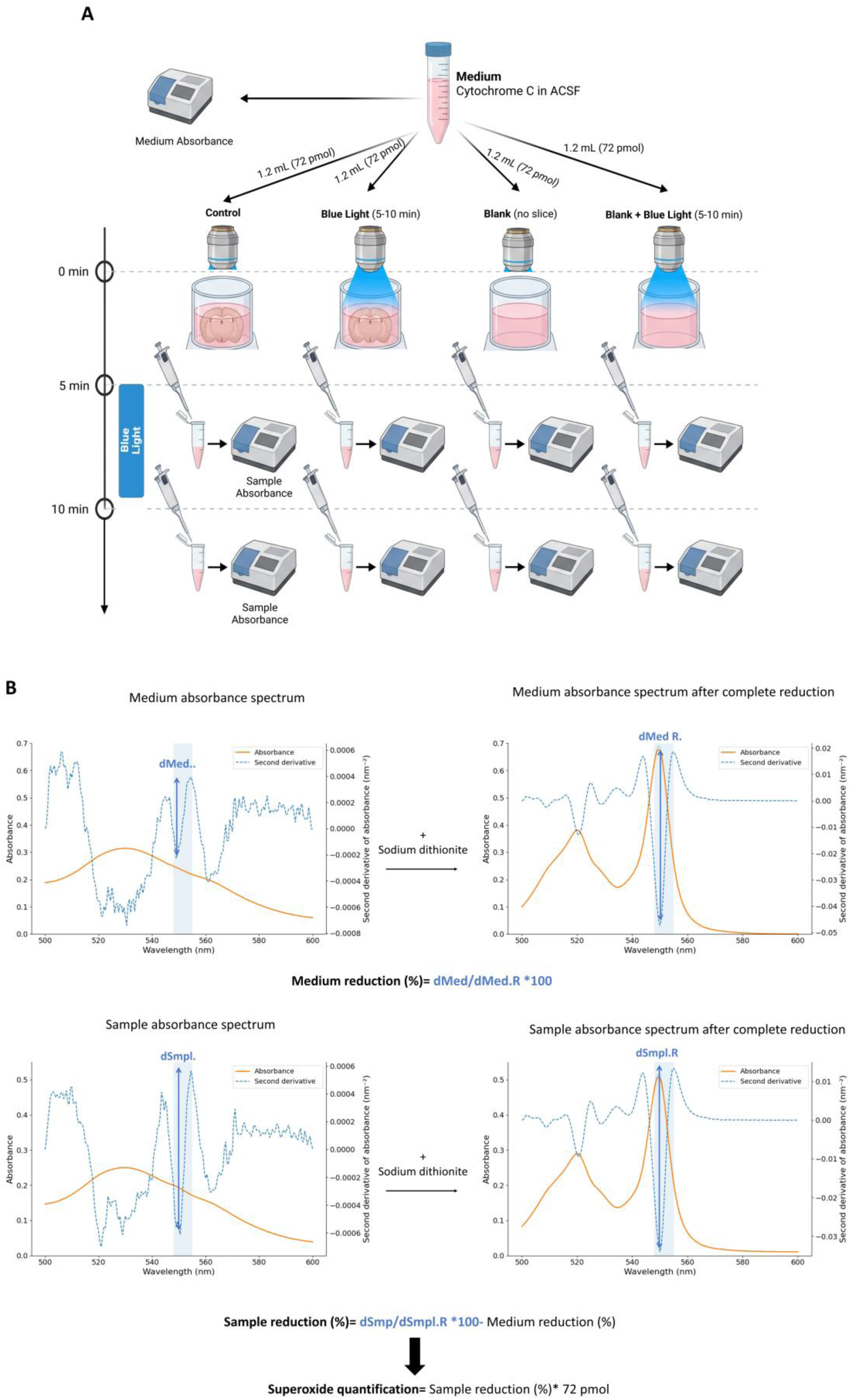
Method for superoxide detection. (A) Schematic representation of the experimental protocol. The recording chamber contained 1.2 ml of oxygenated buffer supplemented with 60 µM acetylated cytochrome c. Samples (200 μL) were collected from the recording chambers at 5 and 10 min after medium injection, either in the presence or absence of cortical slices. In the LED-stimulation groups, blue light was delivered between the two sample collections. (B) Method used to quantify superoxide anions production based on acetylated cytochrome c reduction. The samples were diluted with an equal volume of deionized water, and the amount of reduced acetylated cytochrome c was quantified by measuring the 550-556 nm valley-to-peak amplitude of the second derivative of the absorbance spectrum recorded between 500 and 600 nm. The percentage of acetylated cytochrome c reduction was calculated relative to the complete reduction of the solution induced by sodium dithionite. The baseline percentage of reduced acetylated cytochrome c initially present in the buffer (<1%) was subtracted from that measured in each sample. The amount of acetylated cytochrome c reduced during the experimental procedure was then calculated by multiplying the total amount of acetylated cytochrome c in the recording chamber (72 nmoles) by the baseline-corrected fractional reduction.

**Supplementary Figure 4.**
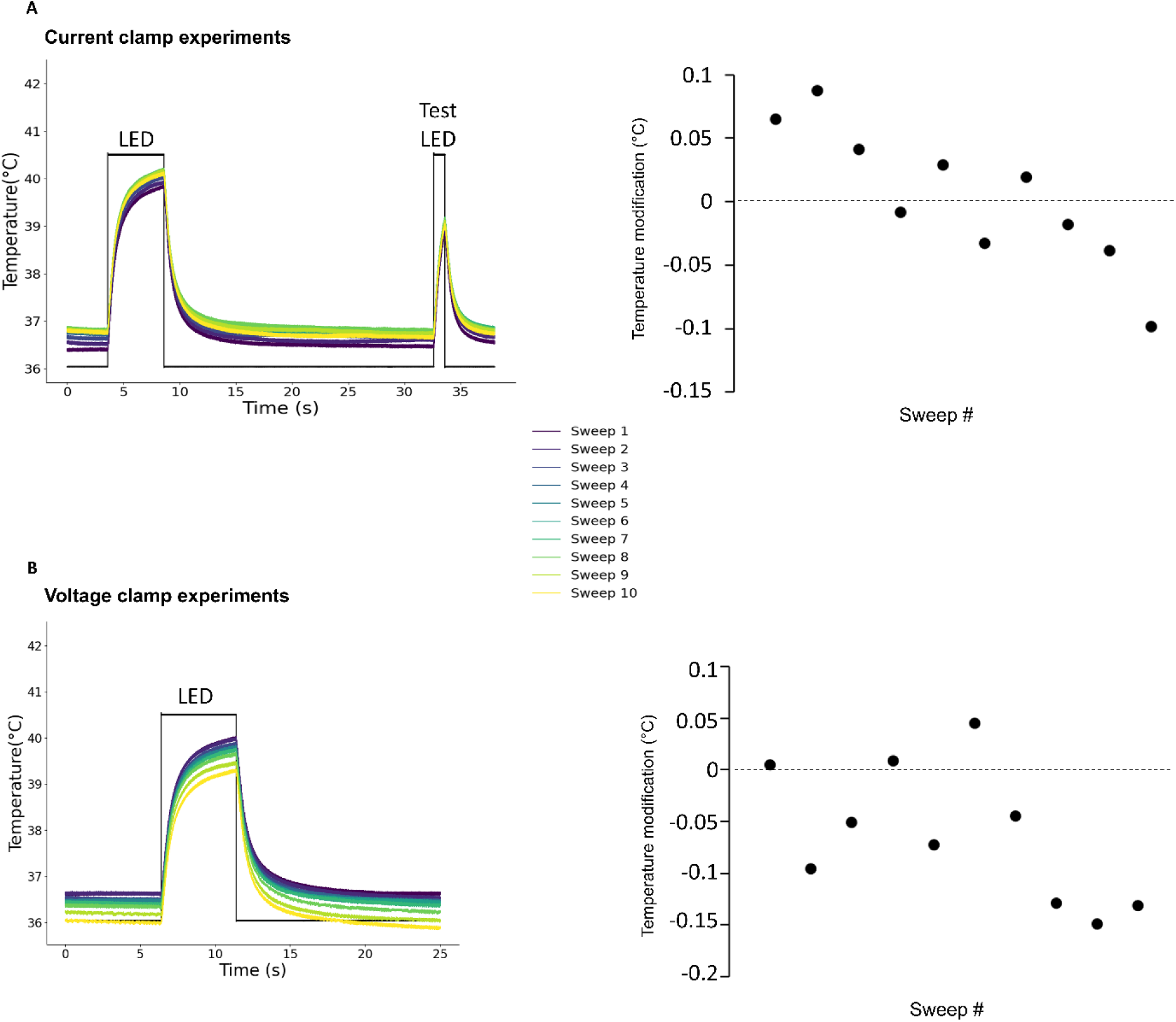
The change in tissue temperature induced by the light returned to baseline within the duration of the sweep. A) Current clamp experiment. Left, time course of tissue temperature over 10 consecutive sweeps. Right, quantification of the absolute temperature difference between the pre-LED and pre-test LED periods, measured during the last 0.5 s preceding each LED stimulus. B) Voltage clamp experiment. Left, time course of tissue temperature over 10 consecutive sweeps. Right, quantification of the absolute temperature difference between the pre-LED period and the last 0.5 s of each sweep.

**Supplementary figure 5.**
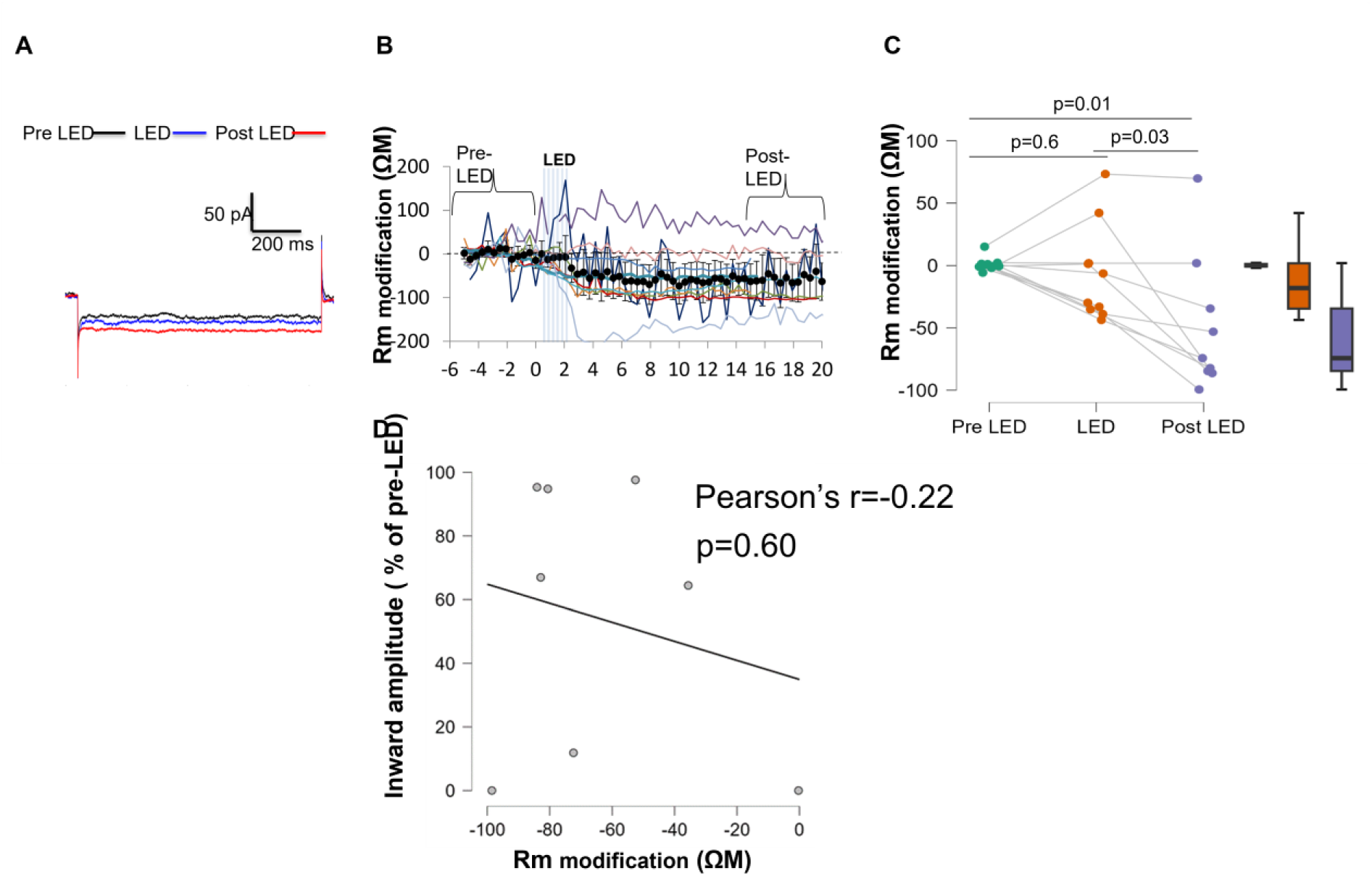
Light stimulation produces a long-lasting decrease of membrane resistance. **(A)** Example of passive membrane currents induced by 5 mV membrane hyperpolarization used to calculate Rm. (**B)** Temporal evolution of the Rm change. Black dots and bars represent mean and 95% CI, respectively. Colored lines represent single cell evolution**. (C)** Statistical quantification of the average Rm changes in the three periods (Friedman test followed by Conover post hoc comparisons). (D) Linear correlation between LED effect on Rm and inward current. Raw data and analysis are available at https://osf.io/pt4br/p

**Supplementary Figure 6.**
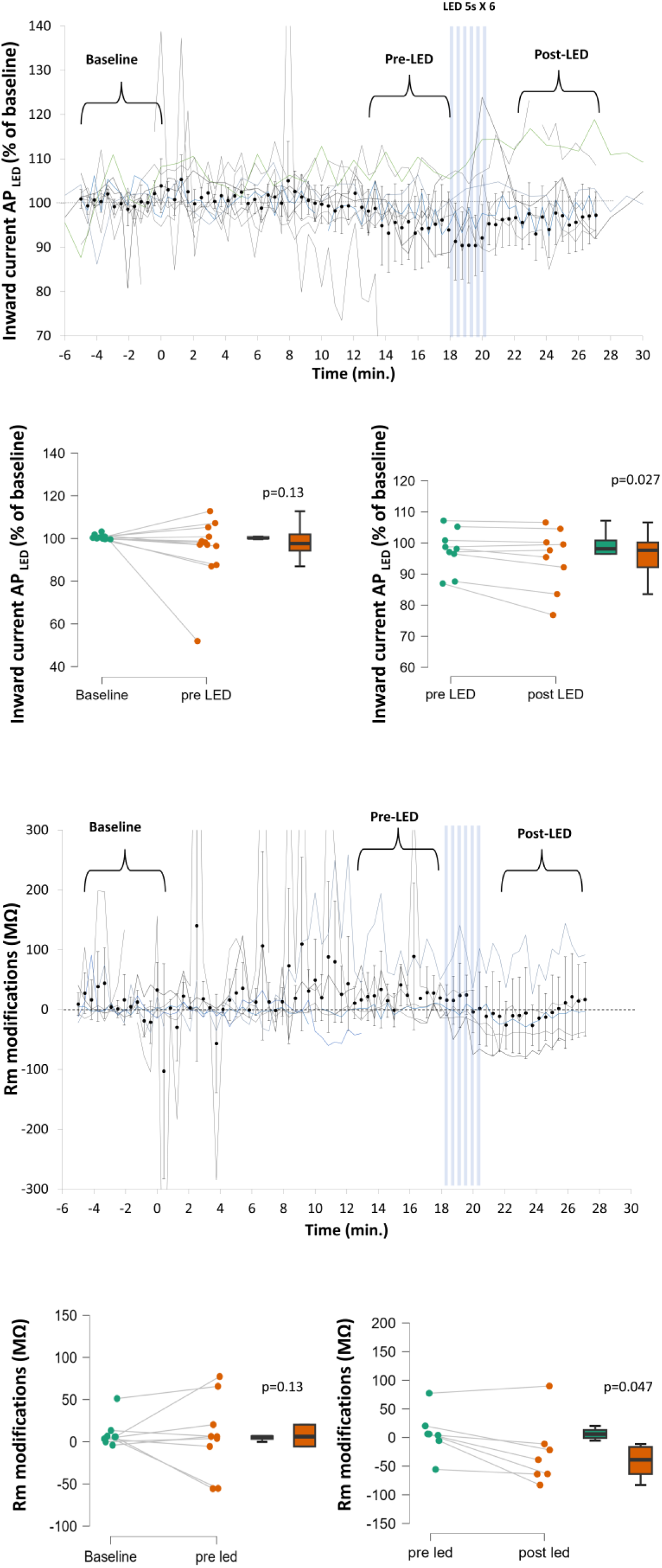
The reduction of inward current and Rm observed in the voltage clamp experiments are not due to spontaneous drift of these parameters. **A)** Inward current evolution. Ai. Time course of inward current activity modification. Aii. Statistical quantification of the spontaneous modification of the inward current. Aiii Statistical quantification of the effect produced by LED stimulation on the inward current. **B)** Rm evolution. Bi. Time course of spontaneous Rm modification. Bii. Statistical quantification of the spontaneous modification of Rm. Aiii Statistical quantification of the effect produced by LED stimulation on Rm. Black dots and bars represent the mean and 95% confidence interval, respectively. Colored lines represent individual cell responses.

**Supplementary figure 7.**
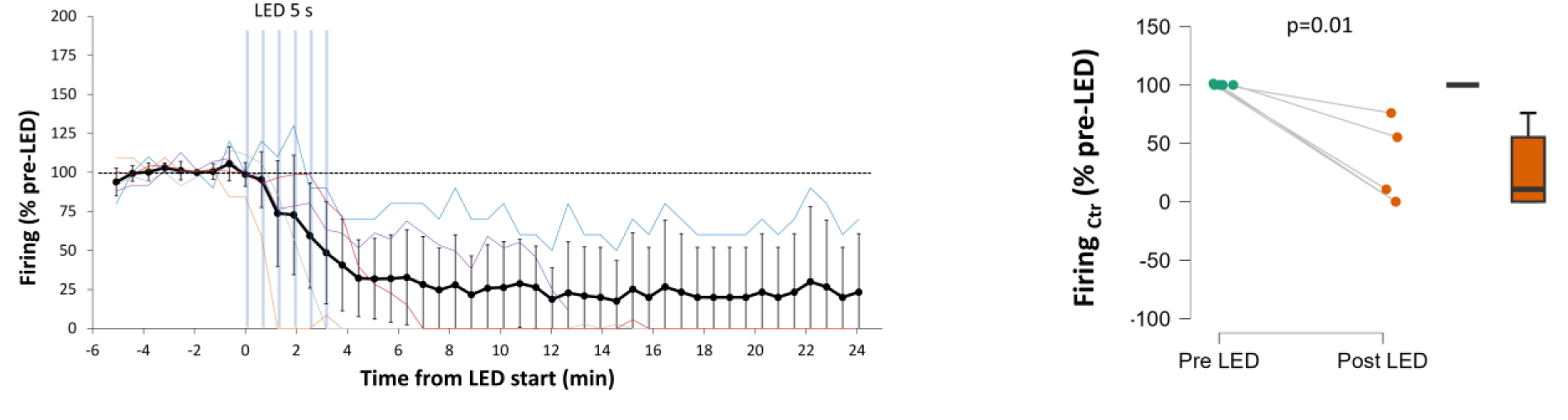
Long recovery time before slicing does not prevent the long-lasting reduction in firing induced by light stimulation in mice. Left, time course of the change in firing rate induced by six LED pulses in neurons from mouse brains kept in cold, oxygenated solution for 20 min before slicing. Right, statistical quantification of the effect of light stimulation on firing rate. Black dots and bars represent the mean and 95% confidence interval, respectively. Colored lines represent individual cell responses.

**Supplementary Figure 8.**
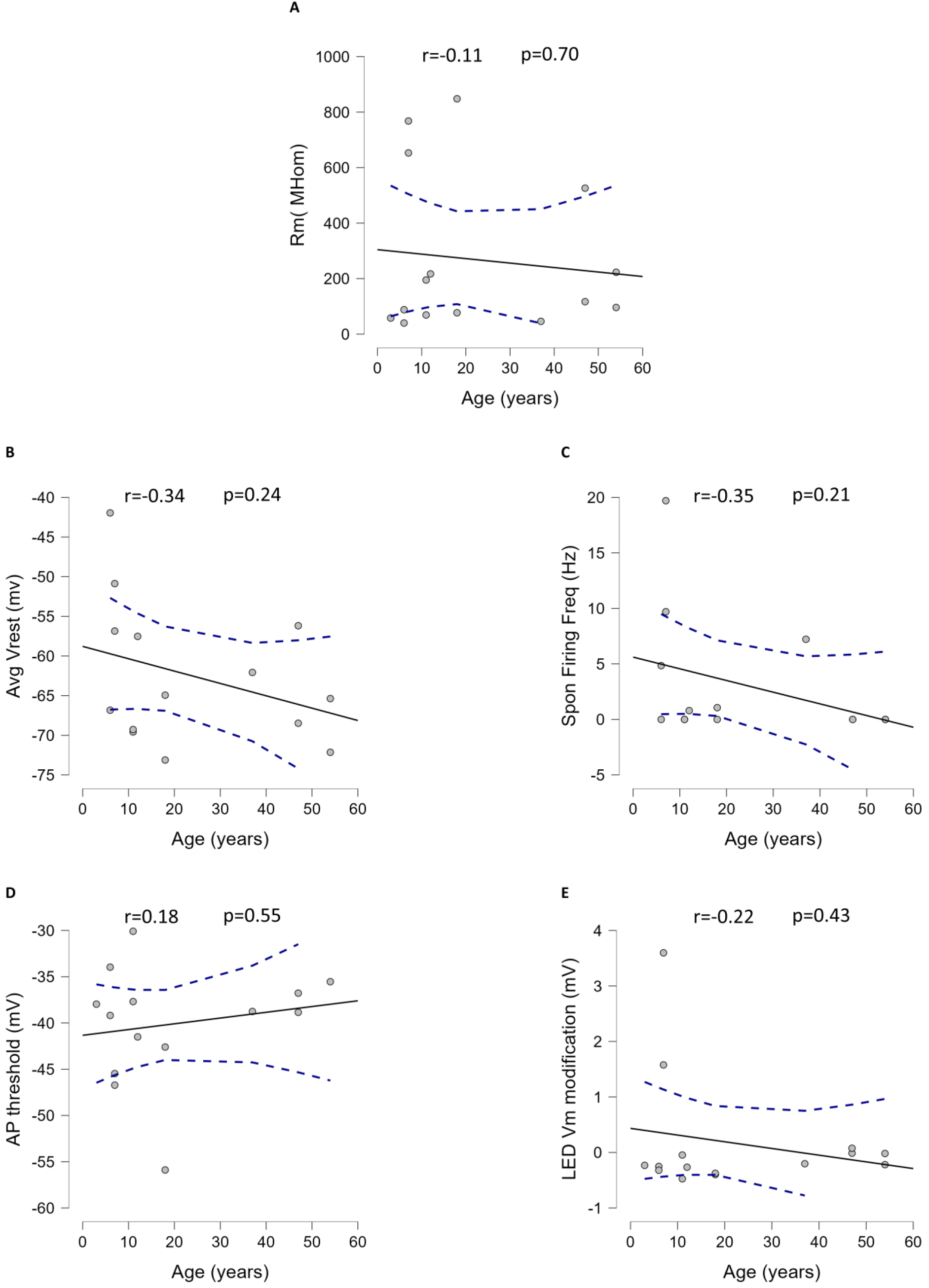
Age of patient as determinant of light effect on human neuronal biophysical properties (A-D). and light modification of membrane potential (E).

**Supplementary Figure 9.**
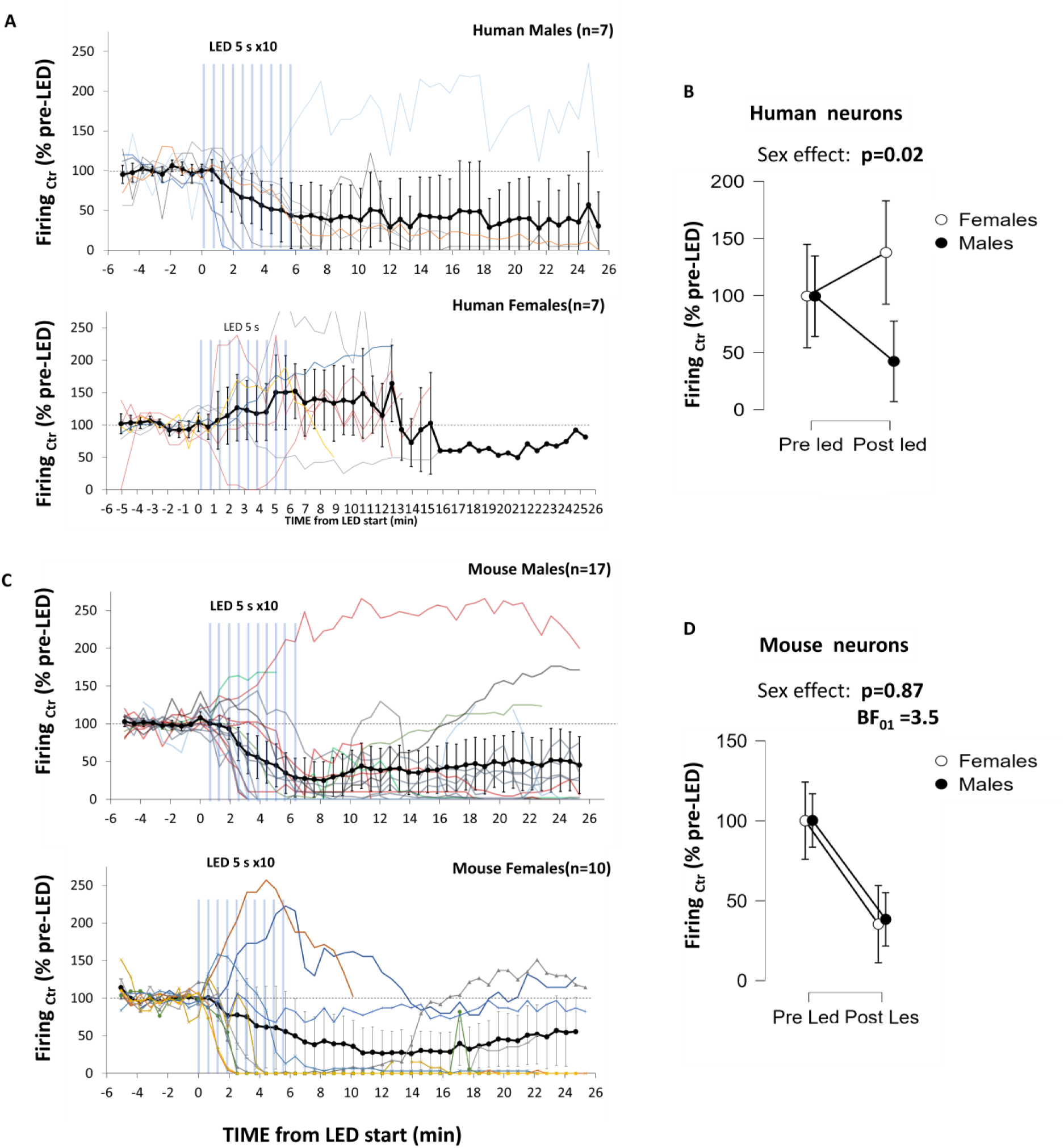
Sex effects on long-lasting modification of firing activity are present only in neurons from humans. **A)** Time course of firing activity modification induced by light stimulation in human neurons. **B)** 2×2 ANOVA of long-lasting light effects in human neurons. **C)** Time course of firing activity modification induced by light stimulation in mouse neurons. **D**) 2×2 ANOVA of long-lasting light effects in mouse neurons. Raw data available at: https://osf.io/2wfy5/

**Supplementary figure 10.**
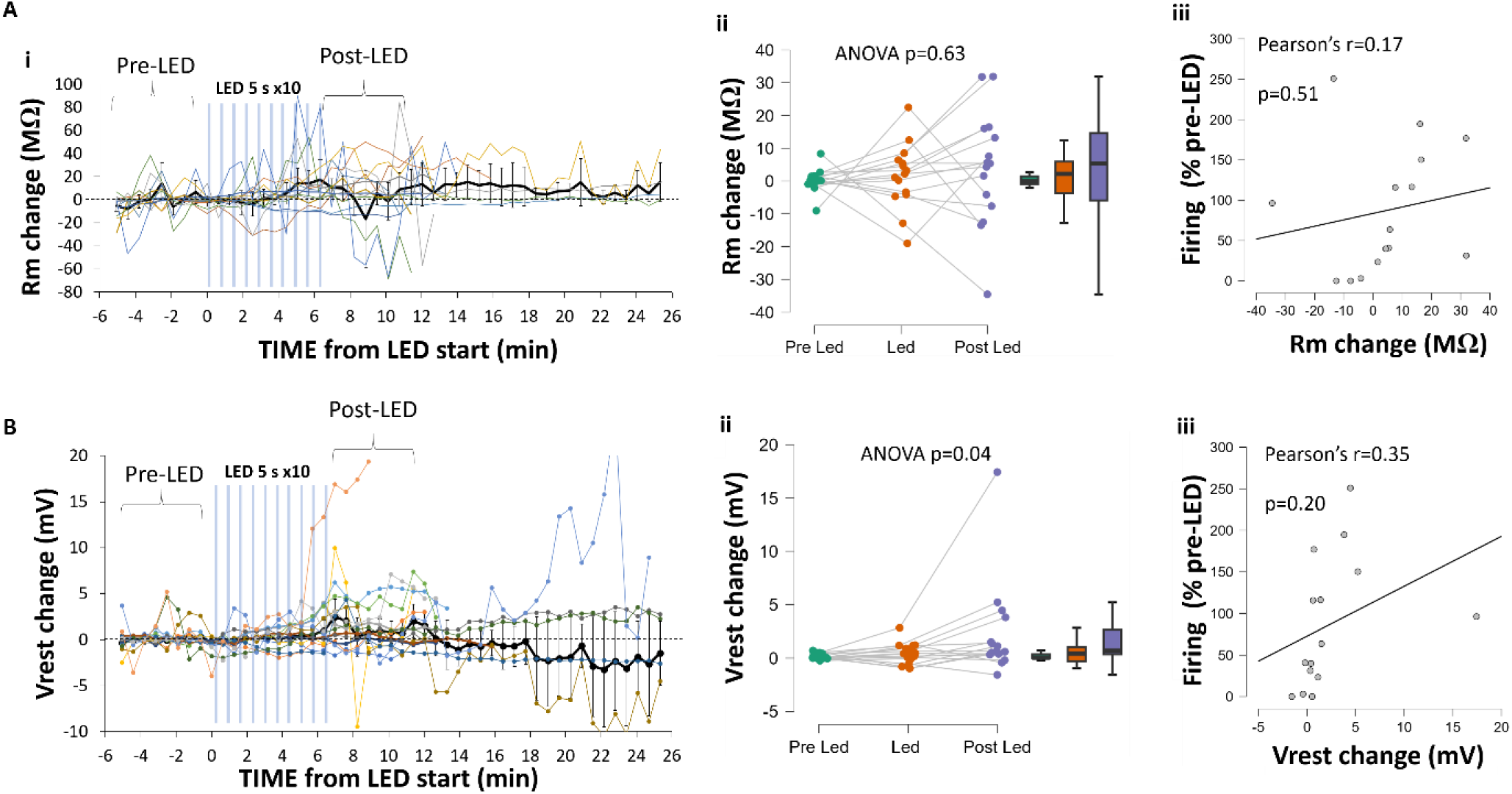
Light effect on membrane resistance and resting membrane potential on human neurons. A) Light effect on membrane resistance. Bii. Time course of the modification of Rm induced by 10 LED pulses. Biii. Statistical quantification of the light effect on Rm across three time periods. B) Light effect on resting membrane potential. Cii. Time course of the modification of Vrest induced by 10 LED pulses. Ciii. Statistical quantification of the light effect on Vrest across three time periods. Black dots and bars represent the mean and 95% confidence interval, respectively. Colored lines represent single cell responses. Raw data available at: https://osf.io/2wfy5/

